# The Identification of IL-4 as a Regulator of Chimeric Antigen Receptor T Cell Exhaustion

**DOI:** 10.1101/2023.09.28.560046

**Authors:** Carli M. Stewart, Elizabeth L. Siegler, R. Leo Sakemura, Michelle J. Cox, Truc Huynh, Brooke Kimball, Long Mai, Ismail Can, Claudia Manriquez Roman, Kun Yun, Olivia Sirpilla, James H. Girsch, Ekene Ogbodo, Wazim Ismail, Alexandre Gaspar Maia, Justin Budka, Jenny Kim, Nathalie Scholler, Mike Mattie, Simone Filosto, Saad S. Kenderian

**Affiliations:** T Cell Engineering, Mayo Clinic, Rochester, MN; Mayo Clinic Graduate School of Biomedical Sciences, Mayo Clinic, Rochester, MN; Department of Molecular Pharmacology and Experimental Therapeutics, Mayo Clinic, Rochester, MN; Division of Hematology, Mayo Clinic, Rochester, MN; Department of Molecular Medicine, Mayo Clinic, Rochester, MN; Department of Lab Medicine and Pathology, Mayo Clinic, Rochester, MN; Department of Oncology, Gilead Sciences Inc., Foster City, CA, USA; Department of Immunology, Mayo Clinic, Rochester, MN

**Author notes:** **Corresponding Author:** Saad Kenderian, M.D., Division of Hematology, Mayo Clinic, 200 First Street S.W., Rochester, MN 55905.

**Keywords:** Chimeric Antigen Receptor, Exhaustion, IL-4, Epigenetic Regulation, ZUMA-1

## Abstract

Durable response to chimeric antigen receptor T (CART) cell therapy remains limited in part due to CART cell exhaustion. We investigated the regulation of CART cell exhaustion with three independent approaches including: a genome-wide CRISPR knockout screen using a validated *in vitro* model for exhaustion, RNA and ATAC sequencing on baseline and chronically stimulated CART cells, and RNA and ATAC sequencing on pre-infusion CART cell products from responders and non-responders in the ZUMA-1 clinical trial. Each of these approaches identified IL-4 as a key regulator of CART cell dysfunction. Further, when CART cells were treated with IL-4, they developed signs of exhaustion, but when CART cells were treated with an IL-4 monoclonal antibody, they showed improved antitumor efficacy and reduced signs of exhaustion in preclinical models. Therefore, our study identified both a novel role for IL-4 on CART cells and the improvement of CART cell therapy through IL-4 neutralization.

**Statement of Significance:** Identifying regulators of CART cell exhaustion will not only enhance the field’s understanding of therapeutic failure, but it will also provide avenues to enhance CART cell efficacy. This study reveals both a novel role for IL-4 in exhaustion and a strategy to improve CART cell activity through IL-4 neutralization.

## Introduction

Chimeric antigen receptor T (CART) cell therapy has evolved as a potentially curative therapy in a subset of patients with hematological malignancies^1^. While CART cell therapy results in impressive overall response rates over 70%, durable response rates remain limited to 30-40%, and most patients relapse within the first year of therapy^2–4^. Several mechanisms of CART cell failure have been identified, including the limited *in vivo* expansion and persistence of CART cells^5,6^.

It has become increasingly evident that T cell exhaustion contributes to CART cell failure in the clinic^7,8^. T cell exhaustion is an epigenetically regulated state of dysfunction that results from chronic stimulation through either the T cell receptor (TCR) in CD8^+^ T cells or through the CAR in CART cells^9–12^. It is characterized by phenotypic, functional, transcriptional, and epigenetic changes. Phenotypic alterations include the upregulation of multiple inhibitory receptors on a cell such as programmed cell death protein 1 (PD-1), T cell immunoglobulin and mucin domain-containing protein 3 (TIM-3), cytotoxic T-lymphocyte associated protein 4 (CTLA-4), and lymphocyte-activation gene 3 (LAG-3)^9^. Functional alterations include a decreased ability to proliferate and to produce effector cytokines such as interleukin (IL)-2 and tumor necrosis factor (TNF)-α, followed by losing the ability to produce interferon gamma (IFN-γ) at later stages^9^. In addition, exhausted T cells experience metabolic changes such as impaired glycolysis^13^. Transcriptional and epigenetic changes include alterations in the activity of several transcription factors including: TCF-7, TOX, T-BET, EOMES, PRDM1, NR4A3, BATF, EGR2, and AP-1 and RUNX family members^14–26^. In an effort to control the epigenetic response to chronic stimulation, several studies have used genetic engineering tools to overexpress or knockout individual molecules. Some examples include the overexpression of the AP-1 family member c-Jun and the deletion of molecules such as the methyltransferase DNMT3A, the inflammatory regulators REGNASE-1 and ROQUIN-1, and the transcription factors PRDM1 and NR4A3^11,25,27,28^. While these approaches have both enhanced the field’s understanding of CART cell exhaustion as well as established a framework to prevent its development, a complete understanding of molecular pathways and potential targets for therapeutic intervention has not been achieved thus far.

It is likely that the occurrence of CART cell exhaustion varies by CAR construct and disease type. For example, independent studies have demonstrated that CD28-costimulated CART cells are more susceptible to a state of exhaustion as compared to 41BB-costimulated CART cells^5,29,30^. In addition, CAR constructs that experience a higher occurrence of tonic signaling have also been positively associated with the development of CART cell exhaustion^29^. Further, current literature supports a higher risk of exhaustion in CART cells used for the treatment of solid tumors due to the added challenges of an immunosuppressive tumor microenvironment^31^.

We focused our studies on the epigenetic regulation of exhaustion in CART cells targeting the CD19 antigen (CART19) with a CD28 costimulatory domain (CART19-28ζ) as a model. To do so, we employed the following three independent strategies: 1) a genome-wide clustered regularly interspaced short palindromic repeat (CRISPR) knockout screen in healthy donor CART19-28ζ cells using an *in vitro* model for exhaustion, 2) RNA and assay for transposase-accessible chromatin (ATAC) sequencing on baseline and chronically stimulated CART19-28ζ cells from healthy donors using an *in vitro* model for exhaustion, and 3) RNA and ATAC sequencing on pre-infusion axicabtagene ciloleucel (axi-cel) products from patient responders and non-responders in the pivotal ZUMA-1 clinical trial that led to the initial FDA approval of axi-cel^32^. Collectively, our independent approaches identified IL-4 as a key regulator of CART cell exhaustion. Subsequently, we performed validation studies to confirm the role of IL-4 on CART cell function using *in vitro* and *in vivo* models, ultimately proposing IL-4 neutralization as a strategy to improve CART cell anti-tumor activity.

## RESULTS

### Establishing an *In Vitro* Model for CART19 Cell Exhaustion

To better understand the development of CART cell exhaustion, we first designed an *in vitro* model to induce exhaustion in CART19 cells generated from healthy donor T cells (Fig. 1A). This model was designed to be both scalable and to focus specifically on the development of exhaustion through chronic stimulation of the T cells. In CD8^+^ T cells, exhaustion has been modeled *in vitro* by the chronic stimulation of the T cells through the T cell receptor (TCR)^9,33^. To model this phenomenon in CART19-28ζ cells, we chronically stimulated the CART cells through the CAR with the addition of fresh CD19^+^ target cells to the culture every other day. When CART19-28ζ cells were chronically stimulated with the CD19^+^ mantle cell lymphoma cell line, JeKo-1, they became progressively dysfunctional as evident by reduced CART cell antigen specific proliferation *in vitro* (Fig. 1B). Additionally, chronically stimulated CART cells exhibited phenotypical and functional signs of exhaustion such as the increased expression of multiple inhibitory receptors and the decreased production of effector cytokines such as IL-2 and TNF-α (Supplementary Fig. S1A-S1D). Furthermore, these cells exhibited reduced polyfunctionality as determined by the number of T cells secreting more than one cytokine (Supplementary Fig. S1E).

**Figure 1.**
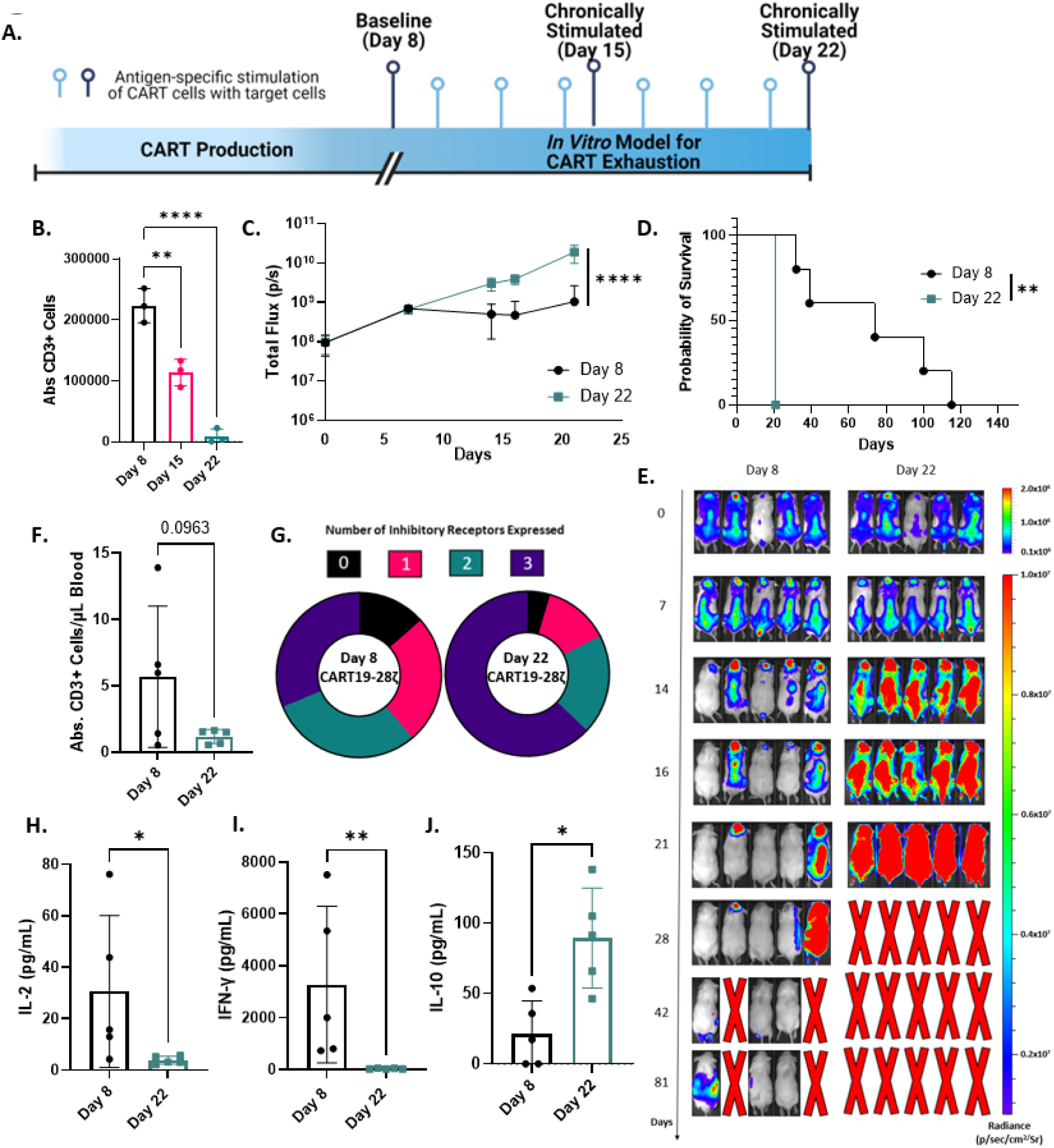
An *in vitro* model for chronic stimulation induces phenotypical and functional signs of exhaustion in CART19-28ζ cells. **A.** Schematic depicting an *in vitro* model for exhaustion in CART cells. **B.** Absolute CD3^+^ cell count, as determined with flow cytometry, after Day 8 (baseline), Day 15 (1-week of chronic stimulation), and Day 22 (2-weeks of chronic stimulation) CART19-28ζ cells were co-cultured with JeKo-1 target cells at a 1:1 ratio for 5 days (One-way ANOVA with three biological replicates, two technical replicates per biological replicate). **C.** *In vivo* antitumor activity of Day 22 and Day 8 CART19-28ζ in a JeKo-1 xenograft model. NOD-*scid* IL2Rgnull (NSG) mice were engrafted with the CD19^+^ luciferase^+^ JeKo-1 cells (1×10^6^ cells I.V.). Mice underwent bioluminescent imaging weekly to confirm engraftment and to monitor tumor burden. Total flux is depicted over time following treatment with CART19-28ζ (0.9×10^6^ cells I.V.) on Day 0 (Two-way ANOVA with n=5 mice per group). **D.** Overall survival curve based on JeKo-1 xenograft mouse model. (Log-rank (Mantle-Cox) test with n=5 mice per group) **E.** Bioluminescent imaging of the tumor growth in the JeKo-1 xenograft mouse model. **F.** *In vivo* CART cell expansion as determined by absolute count of hCD45^+^CD3^+^ cells by flow cytometry per μL of blood on Day 15 of the JeKo-1 xenograft model. (t-test with n=5 mice per group) **G.** Circle graphs showing the average portion of CART cells expressing 0 (black), 1 (pink), 2 (green), or 3 (purple) inhibitory receptors based on flow cytometry detection of PD-1, CTLA-4, and TIM-3 on human CD3^+^ cells in the peripheral blood of mice on day 15 of the JeKo-1 xenograft model. (Average portion from n=5 mice per group) **H-J.** Human cytokine levels for IL-2, IFN-γ, and IL-10 as determined by Multiplex bead assay, in mouse serum collected from peripheral blood on Day 15 of the JeKo-1 xenograft model. (t-test with n=5 mice per group). (*p<0.05, **p<0.01, and ****p<0.0001).

To further examine the function and phenotype of chronically stimulated CART cells, we utilized a mantle cell lymphoma xenograft mouse model that stress tests CART19 cells (Supplementary Fig. S2A). In this model, mice are allowed to develop high disease burden so that standard CART19 doses fail to induce a complete remission of the tumor. Mice were then randomized to receive treatment with either baseline (Day 8) or chronically stimulated (Day 22) CART19-28ζ cells. Treatment with chronically stimulated CART19-28ζ cells resulted in a significant reduction in anti-tumor activity (Fig. 1C and Supplementary Fig. S2B), and overall survival (Fig. 1D-1E and Supplementary Fig. S2C), compared to treatment with baseline CART19-28ζ. In addition, there was a trend for decreased CART expansion (Fig. 1F and Supplementary Fig. S2D) and a significant upregulation of multiple inhibitory receptors (Fig. 1G and Supplementary Fig. S2E-S2F) on CART cells in the peripheral blood of mice treated with chronically stimulated CART19-28ζ cells. Further, peripheral blood cytokine analysis two weeks following CART cell injection showed significantly decreased levels of several effector cytokines (Supplementary Fig. S2G), including IL-2 (Fig. 1H) and IFN-γ (Fig. 1I), in mice treated with chronically stimulated CART cells. However, these mice also showed significantly elevated levels of the inhibitory cytokine IL-10 (Fig. 1J). These findings both corroborate our *in vitro* findings and align with an exhausted phenotype.

Having demonstrated that our *in vitro* model for CART cell exhaustion leads to phenotypic and functional changes associated with T cell exhaustion, we next tested whether this model is applicable to other tumor models and CAR constructs. Consistent with our initial data, chronic stimulation of CART19-28ζ cells with the CD19^+^ acute lymphoblastic leukemia cell line, NALM-6 (Supplementary Fig. S3), or chronic stimulation of CART19-BBζ cells with JeKo-1 cells (Supplementary Fig. S4) resulted in functional and phenotypical signs of T cell exhaustion. As such, our *in vitro* model is a representative model for inducing CART cell exhaustion in different tumor models and different CAR constructs.

### Genome-Wide CRISPR Knockout Screen Identifies the Regulation of IL-4 as a Key Pathway in CART Cell Failure

To investigate genes and pathways that can be altered to protect CART cells from exhaustion, we conducted a genome-wide CRISPR knockout screen using healthy donor CART19-28ζ cells. To do this, we scaled our *in vitro* model for CART cell exhaustion by transducing 100 million CART cells with the GeCKO v2 library A on Day 2 at a multiplicity of infection (MOI) of 0.3 (Fig. 2A)^34^. Then, we selected for transduced cells through treatment with puromycin from Day 3 to Day 8.

**Figure 2.**
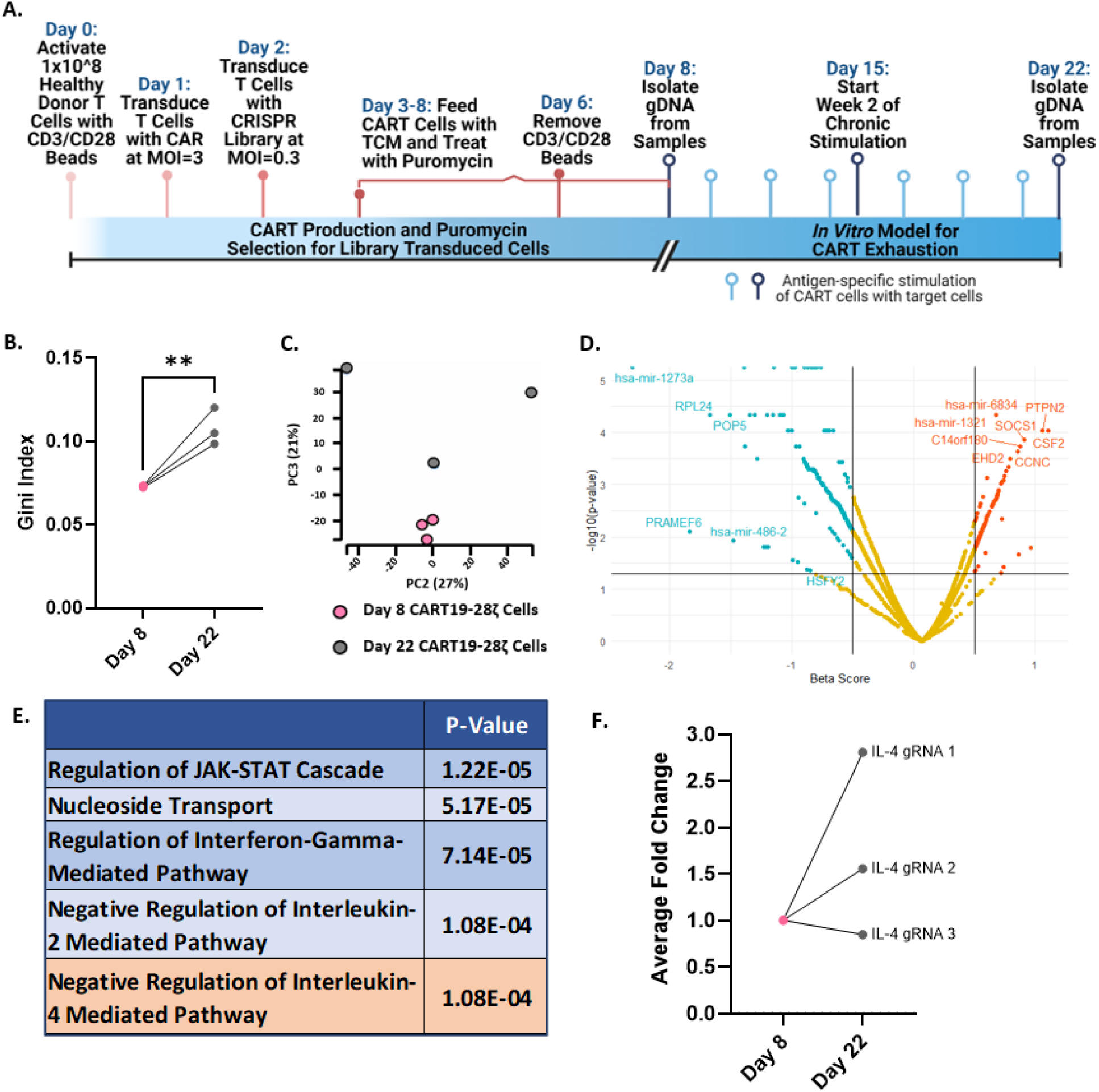
A genome-wide CRISPR knockout screen identifies a role for the IL-4 pathway in the development of CART cell dysfunction resulting from chronic stimulation. **A.** Schematic depicting the genome-wide CRISPR knockout screen conducted in healthy donor CART19-28ζ cells. **B.** The Gini index on Day 8 and Day 22 of the CRISPR screen (Gini index was calculated with MAGeCK-VISPR and compared with a t-test, three biological replicates). **C.** Principal component analysis of gRNA representation in the CRISPR screen at Day 8 and Day 22 (MAGeCK-VISPR analysis with three biological replicates). **D.** Volcano plot showing genes that were positively (red) or negatively (green) selected by Day 22 of the CRISPR screen as compared to Day 8. **E.** Top pathways identified by gene ontology enrichment analysis of the positively selected genes (FDR<0.25) from Day 8 to Day 22 of the CRISPR screen (three biological replicates). **F.** Average fold-change of IL-4 gRNA representation from Day 8 to Day 22 of the CRISPR screen from three biological replicates. (**p<0.01)

By Day 22 of the chronic stimulation assay, we observed positive selection of guide RNAs (gRNAs) as seen by an increase in the Gini index from Day 8 to Day 22 (Fig. 2B) and principal component analysis (PCA) (Fig. 2C). Additionally, top results from gene set enrichment analysis of the negatively selected genes include pathways associated with ribosomal genes and translational processes (Supplementary Fig. S5). This is expected as knockout of these pathways leads to a strong negative selection phenotype^35^. Among the top positively selected genes, there are several genes that have previously been associated with CART cell dysfunction (*CSF2*, *SOCS1*, and *PTPN2*) (Fig. 2D). Knockout of these genes with CRISPR Cas9 in previously published studies has resulted in increased proliferative ability and decreased signs of CART cell dysfunction^36–39^. Together, this data indicates that the genome-wide CRISPR knockout screen effectively identified positively, and negatively selected genes associated with CART cell dysfunction.

In order to investigate key pathways associated with CART cell exhaustion, we performed gene ontology enrichment analysis on the list of positively selected genes. This analysis revealed a role for several cytokine signaling pathways in CART cell exhaustion including the IFN-γ, IL-2, and IL-4 pathways (Fig. 2E). In an independent casual network analysis of the positively selected genes with QIAGEN ingenuity pathway analysis (IPA), the IL-4 receptor (IL4R) was identified as a top regulator (Supplementary Fig. S6). Upon closer investigation of the gRNAs targeting IL-4 in the screen, two out of the three gRNAs targeting IL-4 had enhanced representation in the chronically stimulated (Day 22) samples as compared to baseline (Day 8) samples (Fig. 2F). This suggests that the IL-4 pathway is involved in CART cell exhaustion.

### Transcriptomic and Chromatin Landscape Interrogation of Chronically Stimulated CART Cells Reveals IL-4 as a Regulator of Exhaustion

To further interrogate the top pathways identified with the genome-wide CRISPR knockout screen, and to enhance the field’s understanding of the epigenetic regulation of CART cell exhaustion, we interrogated the transcriptome and chromatin accessibility pattern of baseline and chronically stimulated CART19-28ζ cells from our *in vitro* model for exhaustion. RNA sequencing of baseline (Day 8) and chronically stimulated (Day 15) CART19-28ζ cells revealed the development of a distinct transcriptomic profile (Fig. 3A). Several genes that have previously been correlated with an exhausted phenotype, such as *EOMES*, *IL10RA*, and *HAVCR2* (TIM-3), were confirmed to be upregulated in chronically stimulated CART19-28ζ cells (Fig. 3B). Additionally, the transcription of AP-1 family members that have previously been negatively correlated with the development of exhaustion, such as *FOS* and *FOSL2,* were downregulated (Fig. 3B)^27^.

**Figure 3.**
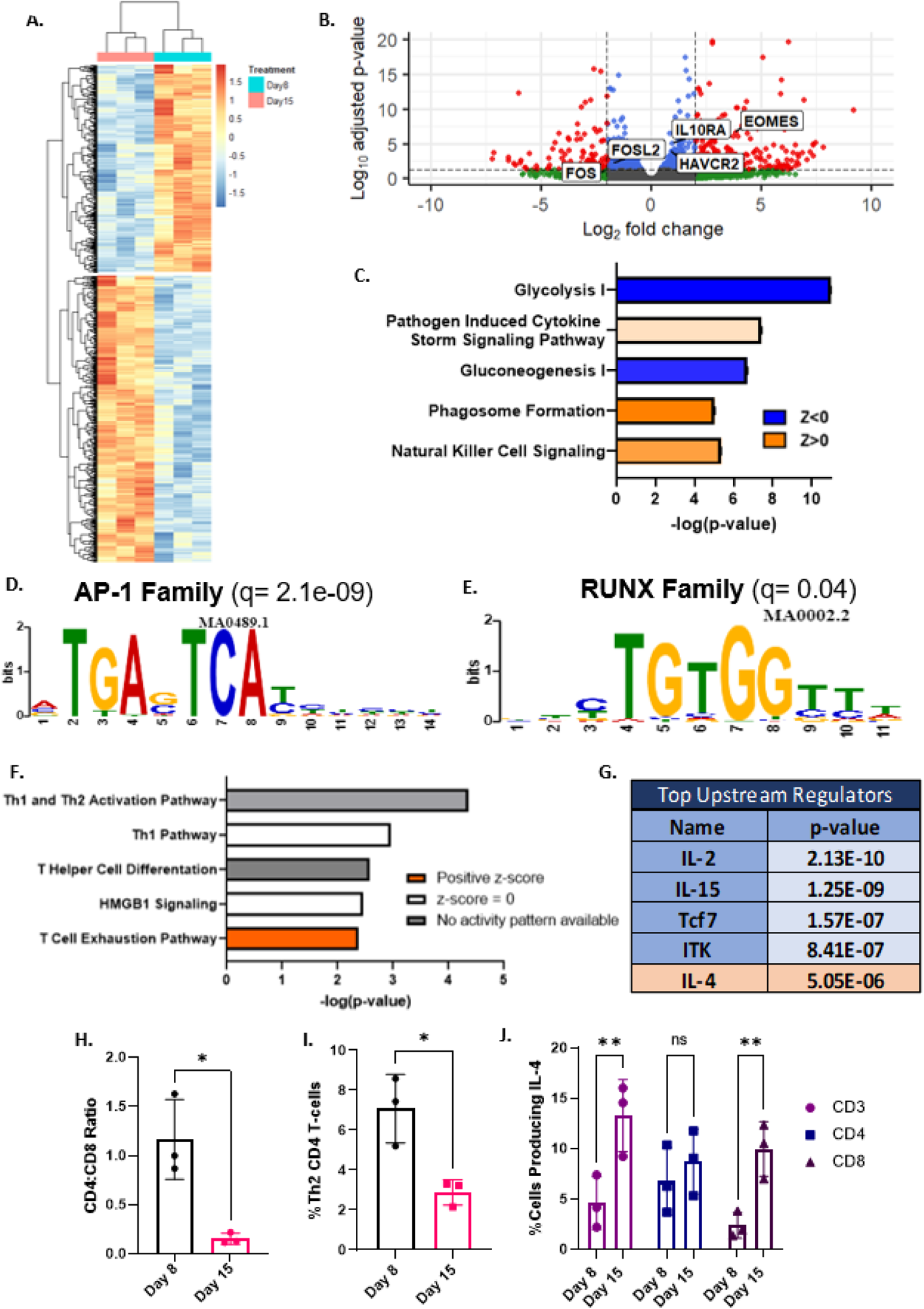
Transcriptomic and chromatin accessibility interrogation reveals a novel role for IL-4 in the development of CART cell exhaustion that is independent of its classical role in Th2 polarization. **A-B.** Heat map and volcano plot showing differentially expressed genes when comparing chronically stimulated (Day 15) to baseline (Day 8) CART19-28ζ cells (DESEQ2 with three biological replicates, padj<0.05). **C.** Top differentially regulated pathways as determined by QIAGEN IPA analysis of differentially expressed genes (padj<0.05). **D-E.** Motifs enriched in chronically stimulated as compared with baseline CART19-28ζ cells (MEME/TOMTOM software with three biological replicates). **F-G.** Top canonical pathways and upstream regulators as identified by QIAGEN IPA analysis of genes that were both differentially expressed (DESEQ2 with three biological replicates, padj<0.05) and differentially accessible (DiffBind with DESEQ2 using three biological replicates, padj<0.05). **H.** Ratio of CD4^+^ to CD8^+^ CART19-28ζ cells in baseline and chronically stimulated cell populations from the *in vitro* model for exhaustion (three biological replicates, two technical replicates per biological replicate, t-test). **I.** The percent of CD4^+^ CART19-28ζ cells that are Th2 polarized as determined by CCR6^-^CCR4^+^CXCR3^-^ in baseline and chronically stimulated CART19-28ζ cell populations (three biological replicates, two technical replicates per biological replicate, t-test). **J.** The percent of either CD3^+^, CD4^+^, or CD8^+^ CART19-28ζ cells producing IL-4 as determined by intracellular staining for IL-4 by flow cytometry following four hours of antigen-specific CAR stimulation through co-culturing Day 15 or Day 8 CART19-28ζ cells with JeKo-1 cells at a 1:5 effector-to-target (E:T) cell ratio (three biological replicates, two technical replicates per biological replicate, 2-Way ANOVA). (*p<0.05, **p<0.01, ns: p>0.05)

To determine the activity of signaling pathways during the development of exhaustion, we performed pathway analysis using both the list of differentially expressed genes and their respective fold changes. This analysis revealed a strong downregulation in pathways associated with metabolism such as glycolysis and gluconeogenesis and highlighted the importance of cytokine signaling in the development of exhaustion (Fig. 3C).

Next, we performed ATAC sequencing on baseline (Day 8) and chronically stimulated (Day 15) CART19-28ζ cells to investigate the epigenetic changes responsible for the development of exhaustion. Consistent with the transcriptomic changes and existing literature, there was enhanced chromatin accessibility at exhaustion-related gene loci (e.g., *PDCD1* and *ENTPD1*) in our chronically stimulated samples (Supplementary Fig. S7A-B) as well as enhanced motif accessibility for AP-1 and RUNX family members (Fig. 3D-3E)^27,33^.

After verifying that known transcriptomic and epigenetic changes associated with exhaustion are seen following chronic stimulation of CART cells in our *in vitro* model for exhaustion, we next asked how the development of exhaustion was being regulated. To uncover epigenetic regulators of CART cell exhaustion, we overlapped the genes that were both differentially expressed and differentially accessible. Then, using QIAGEN IPA, we evaluated top affected pathways and upstream regulators by using both the list of overlapped genes and their respective fold changes, as determined with RNA sequencing. The top enriched pathway was the T cell exhaustion pathway (Fig. 3F). Other pathways with indeterminant enrichment statuses include pathways involved with Th1, Th2 or T helper cell activation or differentiation (Fig. 3F). Top predicted upstream regulators include IL-2, IL-15, TCF-7, ITK, and IL-4 (Fig. 3G). IL-2, IL-15, TCF-7 and ITK have previously been linked to the development of T cell exhaustion^9,16,40–42^. However, while IL-4 has classically been associated with Th2 polarization in CD4^+^ T cells, its role in the development of CART cell exhaustion has not been well studied.

### IL-4 is a Regulator of CART Cell Exhaustion Independent of its Role in Th2 Polarization

Given both the identification of IL-4 as a top upstream regulator in the development of exhaustion and the inclusion of pathways related to T helper cell differentiation in the top affected pathways, we asked whether IL-4 was identified as a top upstream regulator as a result of changes in the T helper cell polarization or as a result of the development of exhaustion.

To approach this question, we utilized multiple independent approaches to differentiate T cell exhaustion from T helper cell polarization in our *in vitro* model for CART cell exhaustion. First, we observed a sharp decrease in the CD4^+^ population of CART19-28ζ cells following chronic stimulation, an established finding during the development of exhaustion (Fig. 3H)^9^. Second, we inspected the Th1 and Th2 population of cells within the declining CD4^+^ population and observed a significant decrease in the Th2 population (Fig. 3I) and an increase in Th1 population (Supplementary Fig. S8A). Third, we evaluated the serum cytokine levels of the Th2 cytokines IL-5 and IL-13 in JeKo-1 xenograft mice treated with either baseline (Day 8) or chronically stimulated (Day 22) CART19-28ζ cells and observed a reduction in the levels of these cytokines in mice treated with chronically stimulated CART19-28ζ cells (Supplementary Fig. S8B-S8C). Fourth, we observed an increase in IL-4 production in CART19-28ζ cells following chronic stimulation with JeKo-1 cells (Fig. 3J). Notably, there was a significant increase in IL-4 production in CD8^+^ CART19-28ζ cells, but not in CD4^+^ CART19-28ζ cells (Fig. 3J).

Overall, IL-4 was identified as a top upstream regulator of exhaustion based on epigenetic changes observed from Day 8 to Day 15 of our *in vitro* model for exhaustion. While this model also produces a shift in the T helper cell population, the shift is towards a Th1 population. This is unexpected given IL-4’s role in polarizing CD4^+^ T cells towards a Th2 phenotype. However, we believe IL-4 is mainly identified as a top upstream regulator as a result of its impact on the CD8^+^ population of CART cells. This is supported by both an increase in the production of IL-4 by CD8^+^ CART cells and a significant reduction in CD4^+^ CART cells following chronic stimulation. As a result, we conclude that IL-4 is playing a regulatory role in the development of CART cell exhaustion that is independent of its classical role in the polarization of CD4^+^ T cells towards a Th2^+^ phenotype.

### IL-4 Transcription and Chromatin Accessibility is Elevated in CART Cell Products from Non-Responders in the ZUMA-1 Clinical Trial

Next, we aimed to determine the significance of our findings in patients treated with CART19-28ζ cells. We interrogated the transcriptome and chromatin accessibility pattern of pre-infusion axi-cel products from responders and non-responders in the pivotal ZUMA-1 clinical trial that led to the initial FDA approval of axi-cel^32^. In this analysis, responders were defined as patients that achieved complete remission as best response while non-responders were defined as patients that experienced stable or progressive disease. There was no observed difference in the percent of T cells expressing CAR between responders and non-responders (Supplementary Fig. S9A).

Interrogation of the transcriptome with RNA sequencing of pre-infusion axi-cel products from 6 responders and 6 non-responders showed clustering of non-responder samples (Fig. 4A). The top upregulated genes in non-responders include IL-4 and CCR3, a chemokine receptor that is known to be induced by IL-4 and IL-2 (Fig. 4B)^43^. Additionally, analysis of the differentially expressed genes with QIAGEN IPA identified IL-4 as one of the upstream regulators, along with other genes such as TNF and STAT3 (Fig. 4C). Interestingly, when we evaluate changes in chromatin accessibility between baseline axi-cel products from responders and non-responders, we see many similarities to the findings observed following chronic stimulation of healthy donor CART19-28ζ cells in our *in vitro* model for exhaustion. Motif analysis revealed an enrichment of motif binding sites for EOMES and PRDM1 (Fig. 4D-4E). Both of these transcription factors have previously been associated with the development of exhaustion^22,23^ Additionally, non-responders showed similar enhancement of chromatin accessibility to exhausted cell populations at exhaustion-related gene loci such as *PDCD1*, *HAVCR2* (TIM-3), *EOMES*, and *IL-10* (Supplementary Fig. S10A-S10D). These findings indicate that epigenetic changes in baseline CART cell products contribute to CART cell exhaustion and failure in the clinic.

**Figure 4.**
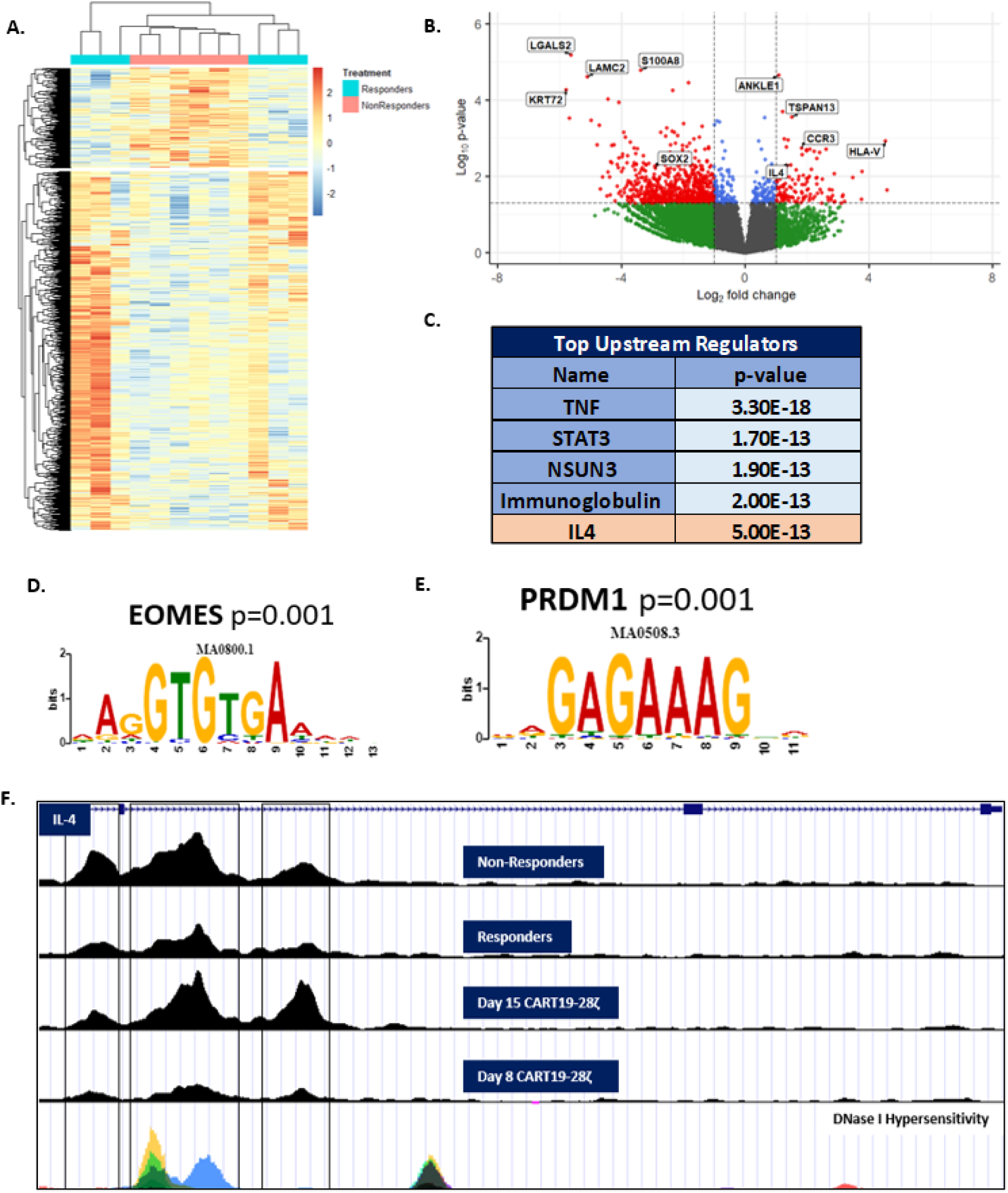
Transcriptomic and chromatin accessibility interrogation of pre-infusion axi-cel products from responders and non-responders in the Zuma-1 clinical trial identifies IL-4 as a regulator of response. **A-B.** Heat map and volcano plot showing differentially expressed genes when comparing pre-infusion axi-cel products from non-responders to products from responders (DESEQ2, six biological replicates per condition, p<0.05). **C.** Top upstream regulators as determined by QIAGEN IPA analysis of differentially expressed genes from non-responder and responder samples. **D-E.** Enriched motifs in baseline products from non-responders as compared to baseline products from responders (MEME/TOMTOM software with six biological replicates per condition). **F.** ATAC signal track of IL-4 gene locus from averaged signal for each experimental condition (axi-cel products from non-responders (n=6), axi-cel products from responders (n=6), Day 15 CART19-28ζ cells (n=3), and Day 8 CART19-28ζ cells (n=3)).

Looking more specifically at the *IL-4* locus, we see enhanced chromatin accessibility in non-responder samples as compared to responders (Fig. 4F). The chromatin accessibility pattern mirrors the change in accessibility observed when healthy donor CART19-28ζ cells are chronically stimulated. In particular, there is enhanced accessibility in both non-responder and chronically stimulated samples at a hypersensitivity site in intron 2 (Fig. 4F). This site correlates with an enhancer locus for IL-4, HS2, that specifically regulates IL-4 without impacting other Th2 cytokines^44^. Collectively, these data indicate that IL-4 regulation in non-responders is consistent with changes seen following the development of CART cell exhaustion in our *in vitro* model for exhaustion.

### Treatment of CART Cells with IL-4 Induces Dysfunction

Due to the identification of IL-4 as a key regulator of CART cell exhaustion through three independent approaches, we performed *in vitro* and *in vivo* studies to directly evaluate phenotypic and functional changes that occur when CART19-28ζ cells are treated with human recombinant IL-4 (hrIL-4). Upon stimulating CART19-28ζ cells once with JeKo-1 target cells in the presence of 20ng/mL hrIL-4 (vs. diluent), CART19-28ζ cells showed signs of dysfunction such as reduced cytotoxicity (Fig. 5A) and a trend for reduced proliferation (Fig. 5B). These changes appear to be independent of a direct impact of hrIL-4 on JeKo-1 cells as treatment of JeKo-1 cells with hrIL-4 alone did not affect their growth or survival (Supplementary Fig. S11A). Further, after three days of co-culturing CART19-28ζ cells with JeKo-1 cells, the concentration of effector cytokines in the supernatant was interrogated with a bead-based multiplex assay. CART19-28ζ cells treated with hrIL-4 secreted lower levels of the effector cytokines TNF-α and IFN-γ as compared with diluent treated cells (Fig. 5C-5D).

**Figure 5.**
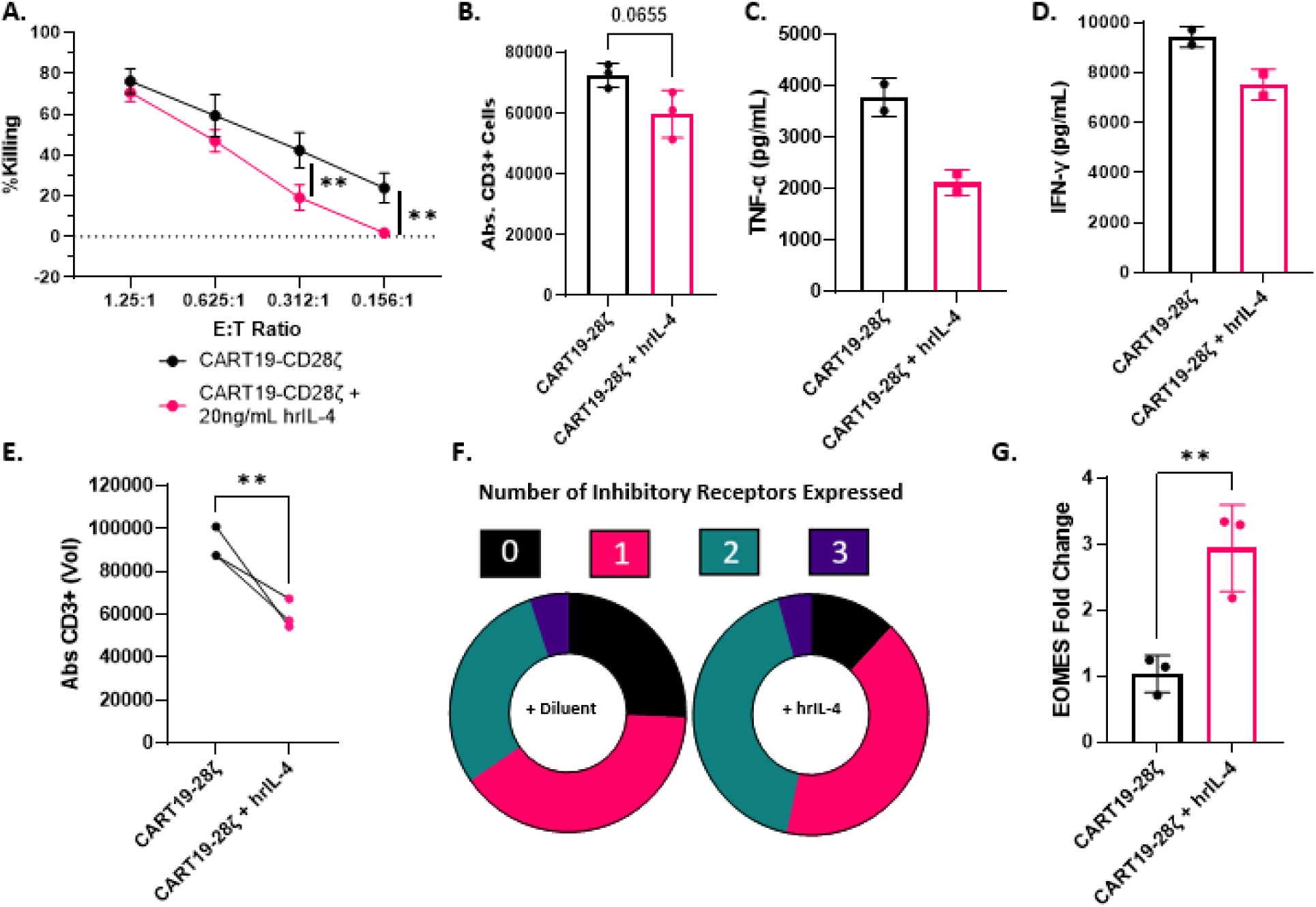
Treatment of CART19-28ζ cells with hrIL-4 leads to phenotypical and functional signs of exhaustion. **A.** Percent killing as measured with bioluminescent imaging after CART19-28ζ cells were co-cultured with luciferase^+^ JeKo-1 cells at various E:T cell ratios for 48 hours in the presence of either 20ng/mL human recombinant IL-4 (hrIL-4) or diluent control (Two-way ANOVA with three biological replicates, two technical replicates per biological replicate). **B.** Absolute CD3^+^ cell count as measured with flow cytometry after CART19-28ζ cells were co-cultured with JeKo-1 cells at a 1:1 E:T cell ratio for five days in the presence of either 20ng/mL hrIL-4 or diluent control (t-test with three biological replicates, two technical replicates per biological replicate). **C-D.** Concentration of TNF-α and IFN-γ in the supernatant as determined with the Milliplex MAP Human High Sensitivity T Cell Panel Premixed 13-plex after CART19-28ζ cells were co-cultured with JeKo-1 cells at a 1:1 E:T cell ratio for three days in the presence of either 20ng/mL hrIL-4 or diluent control (Two biological replicates, two technical replicates per biological replicate). **E.** Absolute CD3^+^ cell count as measured with flow cytometry after Day 15 CART19-28ζ cells that had been chronically stimulated either in the presence of 20ng/mL hrIL-4 or diluent for one week were co-cultured with JeKo-1 cells at a 1:1 E:T cell ratio for five days in the presence of 20ng/mL hrIL-4 (t-test with three biological replicates, two technical replicates per biological replicate). **F.** Circle graph depicting the portion of CART19-28ζ cells expressing 0 (black), 1 (pink), 2 (green), or 3 (purple) inhibitory receptors after being chronically stimulated in the presence of 20ng/mL hrIL-4 or diluent (One representative biological replicate) **G.** The change in the transcription of EOMES as determined with RT-qPCR when CART19-28ζ cells were chronically stimulated for one week in the presence of either hrIL-4 or diluent (t-test with three biological replicates, two technical replicates per biological replicate). (**p<0.01)

Next, we sought to evaluate the impact of IL-4 on CART19-28ζ cells in the presence of chronic stimulation using our *in vitro* model for CART cell exhaustion. Upon chronic stimulation of the CART19-28ζ cells in the presence of 20ng/mL hrIL-4, CART19-28ζ cells showed functional and phenotypic signs of exhaustion such as a reduction in proliferative ability (Fig. 5E) and an increase in the percent of cells expressing multiple inhibitory receptors (Fig. 5F). The increase in the expression of inhibitory receptors appears to be dose-dependent with the greatest increase seen at the highest dose tested, 20ng/mL hrIL-4 (Supplementary Fig. S11B).

With evidence that production of IL-4 is increased in CD8^+^ CART cells upon chronic stimulation and that treatment of CART19-28ζ cells with hrIL-4 causes signs of exhaustion, we next investigated the transcriptional changes responsible for IL-4 induced CART cell dysfunction. Given that 1) a prior study in CD8^+^ T cells showed the induction of the transcription factor EOMES by IL-4^45^ and that 2) EOMES is enriched in both chronically stimulated healthy donor CART cells and non-responder axi-cel products, we asked whether treatment of CART19-28ζ cells with hrIL-4 would induce the expression of EOMES. After chronically stimulating CART19-28ζ cells in the presence of hrIL-4, EOMES transcription is induced as compared to the diluent treated group (Fig. 5G).

Finally, to evaluate if IL-4 induced CART cell dysfunction is dependent on an interaction between the CART19-28ζ cells and target cells, we evaluated both changes in IL-4 production and changes in CART19-28ζ function as a result of hrIL-4 treatment in the presence of irradiated JeKo-1 tumor cells. Following chronic stimulation of CART19-28ζ cells with irradiated JeKo-1 target cells, there is an increase in the percent of CART cells producing IL-4 (Supplementary Fig. 12A). Additionally, upon chronic stimulation of CART19-28ζ cells with irradiated JeKo-1 cells in the presence of hrIL-4, there is an increase in the percent of CART19-28ζ cells expressing multiple inhibitory receptors (Supplementary Fig. S12B-S12C). This suggests that IL-4 induced CART cell exhaustion is due to a direct effect on CART cells and is independent of CART-tumor interactions.

### IL-4 Neutralization Improves the Longevity and Overall Efficacy of CART19-28ζ Cells

Given that IL-4 induces signs of CART cell exhaustion, we next examined whether neutralization of IL-4 with a monoclonal antibody (mAb) could improve CART cell activity. The use of a commercially available IL-4 mAb (clone MP4-25D2), in co-cultures of CART19 and JeKo-1 cells showed complete neutralization of IL-4 (Supplementary Fig. S13A). When CART19-28ζ cells are stimulated with JeKo-1 target cells in the presence of 10μg/mL IL-4 mAb, there is an increase in cytotoxicity (Supplementary Fig. S13B) and an increase in proliferation (Supplementary Fig. S13C) as compared to treatment with an IgG control antibody. Further, when CART19-28ζ cells are chronically stimulated in the presence of 10μg/mL IL-4 mAb, there is a significant decrease in the percent of CART19-28ζ cells that express the inhibitory receptor PD-1 (Supplementary Fig. S13D).

To further investigate the combination of CART19-28ζ cells with an IL-4 mAb, we used our CD19^+^ JeKo-1 stress xenograft mouse model (Fig. 6A), similar to the one depicted in Supplemental Figure 2A. Following CART cell injection, mice were randomized based on tumor burden to weekly intraperitoneal (i.p.) injections of either 10mg/kg IL-4 mAb or an IgG control antibody for 5 weeks. In this model, the majority of mice treated with CART19-28ζ cells and an IgG control antibody were unable to effectively clear their tumor burden (Fig. 6B). Conversely, all mice treated with a combination of CART19-28ζ cells and an IL-4 mAb were able to effectively clear their tumor burden and delay the reappearance of detectable tumor (Fig. 6B). Thus, combining CART19-28ζ cells with an IL-4 mAb resulted in enhanced antitumor activity (Fig. 6C) and a trend for prolonged overall survival (Fig. 6D).

**Figure 6.**
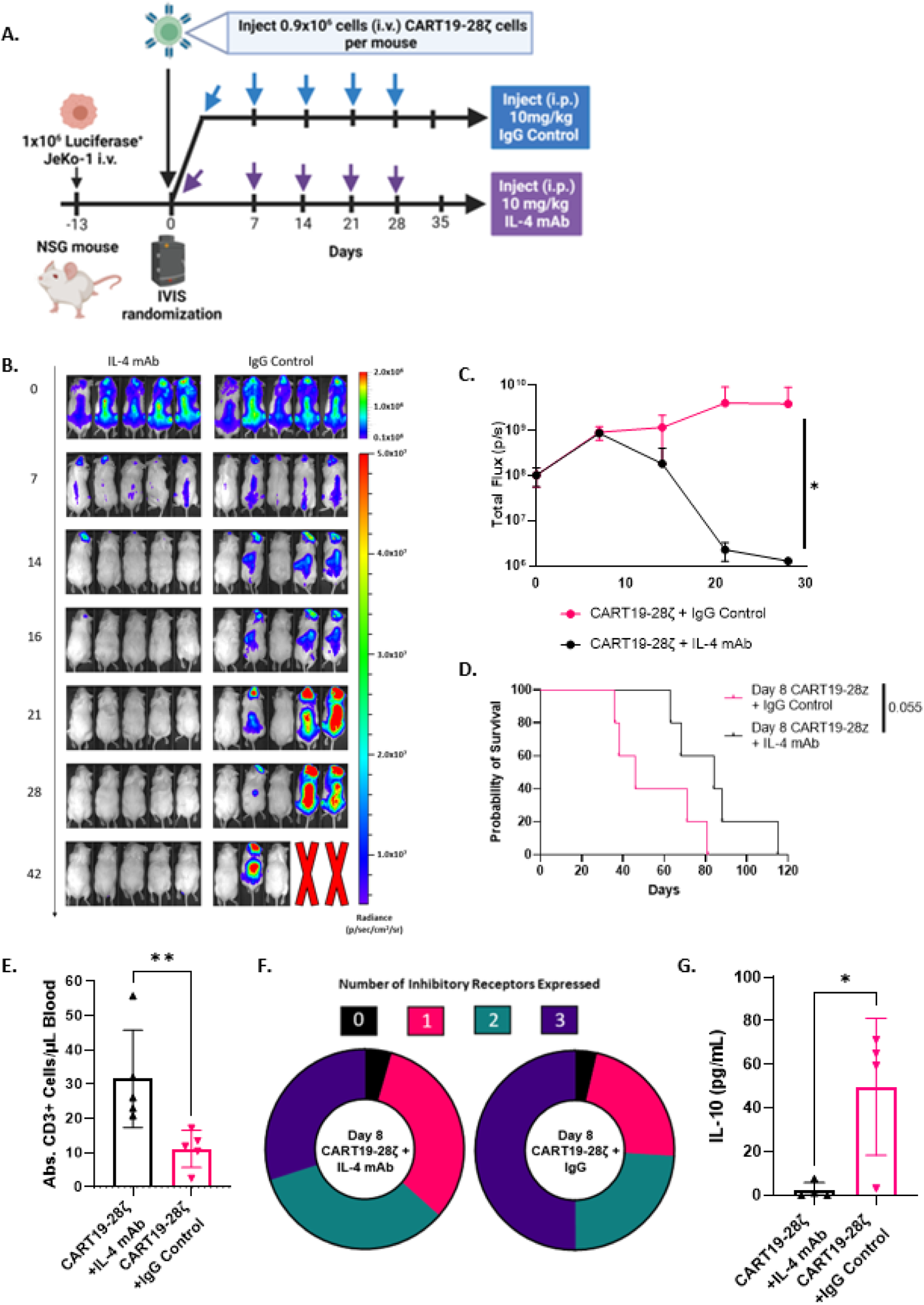
Combination of CART19-28ζ cells with an IL-4 monoclonal antibody improves overall treatment efficacy and CART cell function in an *in vivo* mouse model for mantle cell lymphoma. **A.** Schema for mantle cell lymphoma xenograft mouse model in NSG mice used to test the treatment efficacy of CART19-28ζ cells combined with 10mg/kg IL-4 monoclonal antibody (mAb) as compared with CART19-28ζ combined with 10mg/kg IgG control antibody. **B-C.** Tumor progression as monitored by bioluminescence imaging overtime following injection of CART cells on Day 0 (Two-way ANOVA with n=5 mice per group). **D.** Overall survival curves (Log-rank (Mantle-Cox) test with n=5 mice per group). **E.** Absolute CD3^+^ cells per μL of blood on Day 15 of the *in vivo* study as determined by flow cytometric measurement of cells that are human CD45^+^ and human CD3^+^ after collecting peripheral blood via tail vein bleeding (t-test with n=5 mice per group). **F.** Circle graph showing the average portion of CART cells positive for either 0 (black), 1 (pink), 2 (green), or 3 (purple) inhibitory receptors as determined with flow cytometric detection of human CD3^+^ cells positive for PD-1, TIM-3, and/or CTLA-4 in the peripheral blood of mice on Day 15 of the *in vivo* study (Average value from 5 mice per group). **G.** The concentration of IL-10 in the serum of mice in the *in vivo* model two weeks after the injection of CART cells. Serum was collected through tail bleeding of the mice and cytokine concentration was determined with the use of the Milliplex MAP Human High Sensitivity T Cell Panel Premixed 13-plex (t-test with n=5 mice per group). (*p<0.05 and **p<0.01)

Within this model, we evaluated CART19-28ζ cell function and phenotype through peripheral bleeding two weeks following CART cell injection. The combination of the IL-4 mAb with CART19-28ζ cells improved CART cell expansion (Fig. 6E), reduced the percent of CART cells expressing multiple inhibitory receptors (Fig. 6F), and reduced the secretion of the inhibitory cytokine IL-10 (Fig. 6G and Supplementary Figure S14A). Consistent results were observed in a low-tumor burden CD19^+^ JeKo-1 xenograft mouse model, where all mice were able to clear tumor load regardless of combination therapy (Supplementary Fig. S14B-S14D). Together, our data demonstrates that IL-4 neutralization with a monoclonal antibody reduces exhaustion and improves anti-tumor activity of CART19-28ζ cells.

## DISCUSSION

Our study started with the goal of investigating the development of CART cell exhaustion due to its known role in response to CART cell therapy. To begin our studies, we developed and utilized an *in vitro* model for CART cell exhaustion that resulted in phenotypic, functional, transcriptional, and epigenetic changes associated with exhaustion. Using this *in vitro* model for exhaustion, we performed a genome-wide CRISPR knockout screen that helped us determine genes and pathways that can be altered to protect CART cells from exhaustion, including the IL-4 pathway. In a second approach, we identified IL-4 as a top upstream regulator of CART cell exhaustion by performing RNA and ATAC sequencing on chronically stimulated and baseline CART19-28ζ cells. Finally, in a third approach, we evaluated clinically relevant determinants of CART cell response by performing RNA and ATAC sequencing on pre-infusion axi-cel products from responders and non-responders in the pivotal ZUMA-1 clinical trial. This independent approach not only showed epigenetic changes associated with exhaustion in baseline CART cells from non-responders, but it also identified IL-4 as a top regulator of CART cell dysfunction. Thus, three independent approaches highlighted the involvement of IL-4 in CART cell exhaustion using both preclinical models and clinical trial samples.

While IL-4 has classically been known to promote the polarization of CD4^+^ helper T cells towards a Th2 phenotype, the finding of a regulatory role for IL-4 in CART cell exhaustion is novel^46^. From our studies, we not only observe a sharp decrease in the CD4^+^ population of CART19-28ζ cells following chronic stimulation, but we also see an increase in IL-4 production by CD8^+^, but not CD4^+^ CART19-28ζ cells. This is paired with a decrease in Th2 CD4^+^ CART19-28ζ cells. As such, we believe that this new role for IL-4 is independent of Th2 polarization and is likely due to an effect of IL-4 on CD8^+^ CART19-28ζ cells. However, future studies will need to directly treat CD8^+^ CART19-28ζ cells with hrIL-4 to confirm this hypothesis.

After verifying that treatment of CART cells with hrIL-4 causes signs of exhaustion, we then asked if the development of CART cell exhaustion could be mitigated with IL-4 neutralization. IL-4 neutralization with a monoclonal antibody resulted in improved CART cell antitumor efficacy and expansion both *in vitro* and *in vivo* while also reducing signs of exhaustion such as the expression of multiple inhibitory receptors. These data present a promising therapeutic strategy to improve response to CART cell therapy through combination with an IL-4 monoclonal antibody.

This strategy is potentially translatable to the clinic, as IL-4 targeted therapies are in various stages of clinical development and are generally well tolerated. The IL-4 monoclonal antibody, pascolizumab, was well-tolerated in a phase II clinical trial for the treatment of asthma^47^; dupilumab, a monoclonal antibody blocking IL-4 and IL-13, is FDA-approved for multiple allergic diseases, including eczema and asthma^48^; and pitrakinra, a small molecule inhibitor of IL-4 signaling, has been shown to be safe in a phase II clinical trial for the treatment of asthma^49^. The combination of FDA approved CART19 cells with existing therapeutics to neutralize or antagonize IL-4 is an attractive approach because it avoids further editing of CART cells which results in additional risks and complicates the already timely and expensive CART production process.

While this work focused on the exhaustion of CART19 cells in multiple models of leukemia and lymphoma, the discovery of IL-4 as a regulator of exhaustion is likely applicable beyond hematological malignancies. CART cell therapy has largely failed to gain traction in solid tumor settings due in large part to T cell exhaustion induced by the immunosuppressive tumor microenvironment^31^. IL-4 has been shown to be upregulated in the tumor microenvironment, promote the progression of solid tumors, enhance the generation of immunosuppressive myeloid cells, and dampen antitumor immune responses^50^. Thus, combination of CART cell therapy with and IL-4 neutralizing antibody in the treatment of solid tumors holds the potential to not only directly improve the activity of the CART cells, but also to improve the tumor microenvironment to promote tumor killing.

In summary, our findings utilized relevant preclinical models as well as patient samples from a clinical trial to identify a novel role for IL-4 in CART cell exhaustion that is independent from Th2 polarization. This study also presents IL-4 neutralization as a clinically feasible strategy to prevent CART exhaustion and to enhance CART antitumor efficacy. CART intrinsic dysfunction remains a major challenge to clinical efficacy in both hematological and solid tumors. This study illuminates a novel mechanism for preventing CART exhaustion and a strategy to develop more effective CART cell therapies.

## METHODS

Detailed methods are available in the supplemental materials.

## DATA AVAILABILITY STATEMENT

All data, methods, and study materials will be made available to other researchers upon request. The sequencing data generated in this study will be publicly available in the Gene Expression Omnibus.

## Author Contributions

CMS and SSK conceptualized the project and designed experiments; CMS, MJC, RLS, TH, BK, LM, and ELS performed experiments; CMS analyzed data and prepared manuscript figures; CMS, ELS, and SSK wrote the manuscript; all authors edited and approved the final manuscript; CMS, MJC, and JB performed the bioinformatic analyses, with consultation from WMI and AGM.

## Acknowledgements

CMS, CMR, KY, OS, and JHG are supported by the Mayo Clinic Graduate School of Biomedical Sciences. Schematics are created with BioRender.com.

