## Supplementary material for "The Identification of IL-4 as a Regulator of Chimeric Antigen Receptor T Cell Exhaustion": Methods and Supplemental Figures

**Affiliations:**

**METHODS:**

**Cell Lines:** The mantle cell lymphoma cell line, JeKo-1, and the acute lymphoblastic leukemia cell line, NALM6, were purchased from ATCC, Manassas, VA, USA. For cytotoxicity and *in vivo* experiments, JeKo-1 cells were transduced with luciferase-ZsGreen lentivirus (Addgene, Cambridge, MA, USA) and sorted to 100% purity prior to use. All cell lines were cultured in Roswell Park Memorial Institute (RPMI) 1640 medium (Gibco, Gaithersburg, MD, USA) with 1% Penicillin-Streptomycin-Glutamine (Gibco, Gaithersburg, MD, USA) and either 10% fetal bovine serum (FBS, Sigma, St. Louis, MO, USA) for NALM6 or 20% FBS for JeKo-1 cells. These cell lines were also tested monthly to confirm negative mycoplasma contamination.

**Multi-Parametric Flow Cytometry:** Antibodies were purchased from BioLegend, eBioscience, or BD Biosciences. The preparation of samples for flow cytometry was previously described^1,2^. Flow cytometry was performed with a three-laser CytoFLEX (Beckman Coulter, Chaska, MN, USA). Analyses were performed using Kaluza 2.1 software. More specifically, we performed the following:

*CAR Detection:* Protein L staining with a Biotin Protein L primary antibody (cat. #M00097, GenScript, Piscataway, NJ, USA) and a secondary antibody for streptavidin (cat. #405203, BioLegend, San Diego, CA, USA) or an anti-Whitlow linker antibody was used to detect CAR-28ζ expression on the surface of T cells. A goat anti-mouse F(ab’) antibody (Invitrogen, Carlsbad, CA, USA) was used to detect CAR-41BBζ expression.

*Intracellular Staining of Cytokines:* A degranulation and intracellular cytokine assay was used to determine changes in the production of cytokines as previously described ^1-3^. Briefly, CART cells were co-cultured with JeKo-1 target cells at a 1:5 effector-to-target cell ratio for four hours with the addition of CD28 (clone L293, cat #348040, BD Biosciences, San Diego, CA, USA), CD49d (clone L25, cat #340976, BD Biosciences, San Diego, CA, USA), monesin (cat #420701, BioLegend, San Diego, CA, USA), and CD107a (clone H4A3) FITC (cat #555800, BD Biosciences, San Diego, CA, USA). After four hours of co-culture, intracellular staining of T cells was performed by first staining with Live/Dead Aqua (cat #L34966, Invitrogen, Carlsbad, CA, USA) before using the FIX & PERM^TM^ Cell Permeabilization Kit (cat #GAS001S100 and cat #GAS002S100, Life Technologies, Oslo, Norway). Then, intracellular staining was performed with the following antibodies: IL-2 (clone 5344.111) PE-CF594 (cat #562384, BD Biosciences, San Diego, CA, USA), IL-4 (clone MP4-25D2) APC (cat #554486, BD Biosciences, San Diego, CA, USA), IFN-γ (clone 4S.B3) APC-efluor 780 (cat #47-7319-42, eBioScience, San Diego, CA, USA), GM-CSF (clone BVD2-21C11) BV421 (cat #562930, BD Biosciences, San Diego, CA, USA), and TNF-α (clone Mab11) AF700 (cat #502928, BioLegend, San Diego, CA, USA).

*Surface Staining for Inhibitory Receptors:* To determine changes in the expression of inhibitory receptors from *in vitro* assays, cells were stained with the following antibodies: CD3 (clone SK7) APC-Cy7 (cat #344818, BioLegend, San Diego, CA, USA), CD8 (clone SK1) PerCP (cat #344708, BioLegend, San Diego, CA, USA), PD-1 (clone EH12.2H7) BV-421 (cat #329920, BioLegend, San Diego, CA, USA), TIM-3 (clone F38-2E2) PE (cat #345006, BioLegend, San Diego, CA, USA), CTLA-4 (clone BNI3) PE-Cy7 (cat #369614, BioLegend, San Diego, CA, USA), and LAG-3 (clone 3DS223H) FITC (cat #11-2239-42, eBioscience, San Diego, CA, USA).

*Evaluating Changes in the Th1/Th2 Phenotype by flow cytometry:* To determine changes in the Th1/Th2 phenotype of the CART cells, cells were stained with the following antibodies: CD3 (clone SK7) BV605 (cat #344836, BioLegend, San Diego, CA, USA), CD4 (clone OKT4) FITC (cat #11-0048-42, eBioscience, San Diego, CA, USA), CCR4 (clone L291H4) PerCP (cat #L291H4, BioLegend, San Diego, CA, USA), CCR6 (clone 11A9) APC (cat #560619, BD Biosciences, San Diego, CA, USA), and CXCR3 (clone G02H7) APC-Cy7 (cat #353722, BioLegend, San Diego, CA, USA). Th2 cells were identified as CD3^+^CD4^+^CCR6^-^CCR4^+^CXCR3^-^ and Th1 cells were identified as CD3^+^CD4^+^CCR6^-^CCR4^-^CXCR3^+^.

**Healthy Donor CART Cell Production for *In Vitro* and *In Vivo* Studies:** The use of recombinant DNA in the laboratory was approved by the Mayo Clinic Institutional Biosafety Committee (IBC), IBC number HIP00000252.43. Healthy donor CART cells used in *in vitro* and *in vivo* validation studies were generated as previously reported^1-5^ and as shown in Supplementary Figure S15. Briefly, peripheral blood mononuclear cells (PBMCs) were isolated from de-identified healthy donor blood samples that were obtained under a Mayo Clinic IRB approved protocol by using SepMate PBMC Isolation Tubes (STEMCELL Technologies, Vancouver, Canada). Next, T cells were isolated from the PBMCs through negative selection with an EasySep Human T Cell Isolation Kit with a RoboSep machine (STEMCELL Technologies, Vancouver, Canada). Then, the T cells were activated through the addition of CTS^TM^ (Cell Therapy Systems) Dynabeads^TM^ CD3/CD28 beads (Life Technologies, Oslo, Norway) at a 3:1 bead-to-cell ratio. After 24 hours of bead stimulation, T cells were transduced with lentiviral particles encoding the specific CAR at a multiplicity of infection (MOI) of 3. The generation of lentiviral particles was previously described^1^. The studies included in this paper utilized the following CARs: 1) a second-generation CAR targeting CD19 through an scFv derived from an anti-human CD19 antibody clone FMC63 with a CD28 co-stimulatory domain (CART19-28ζ) and 2) a second-generation CAR targeting CD19 through an scFv derived from an anti-human CD19 antibody clone FMC63 with a 4-1BB co-stimulatory domain (CART19-BBζ)^1^. CART cells were maintained at a concentration of 1x10^6^ cells/mL throughout the production period with T cell media made with X-Vivo15 (Lonza, Walkersville, MD, USA) supplemented with 10% human serum albumin (Corning, NY, USA) and 1% Penicillin-Streptomycin-Glutamine (Gibco, Gaithersburg, MD, USA). On Day 6 of the production period, the CD3/CD28 beads were removed from the cell suspension using magnetic separation before CAR expression was evaluated via flow cytometry as described above. Then, the CART cells were rested in T cell media until Day 8. On Day 8, CART cells were either cryopreserved for future experiments or used fresh for experiments, as indicated. Before use in functional experiments, the cells were thawed in T cell medium and rested overnight.

**CART Cell Production for RNA and ATAC Sequencing:** Healthy donor and pre-infusion patient CART cells used for RNA and ATAC sequencing were generated by KITE Pharma as previously described for the axi-cel clinical product^6,7^, and shipped to the Kenderian Laboratory at Mayo Clinic.

***In Vitro* Model for CART Cell Exhaustion:** Following CART cell production, baseline (Day 8) CART cells were co-cultured with target cells at a 1:1 effector-to-target cell ratio. Several forms of antigen-specific stimulation of CART19 cells were tested and validated including JeKo-1 cells and NALM6 target cells (Fig. 1 and Supplementary Fig. S1-S3). Every other day for a week, CART cells were restimulated with target cells by adding the same number of target cells that was added to baseline CART cells (Fig. 1A). Following one week of chronic stimulation (Day 15), CART cells were isolated through the combined use of CD4 and CD8 microbeads (cat #130-045-101 and cat #130-045-201, Miltenyi Biotec, Auburn, CA, USA). Briefly, CD4 and CD8 beads were added to the washed cell pellet at a 1:1 ratio according to product instructions. Following magnetic separation with LS columns (cat #130-042-401, Miltenyi Biotec, Auburn, CA, USA), purity was verified through staining with anti-human CD3 (clone SK7) APC-Cy7 (cat #344818, BioLegend, San Diego, CA, USA) and Live/Dead Aqua (cat #L34966, Invitrogen, Carlsbad, CA, USA). Then, the function and phenotype of the CART cells was interrogated by evaluating inhibitory receptor expression, cytokine production, and proliferative ability. The remaining Day 15 CART cells were further chronically stimulated by continuing the *in vitro* model for exhaustion for an additional week to generate two-week chronically stimulated CART cells (Day 22 cells). During this additional week, CART cells were re-stimulated every other day with the same number of target cells that was added on Day 15. Bead separation of the CART cells from the co-culture did not impair the activity of the CART cells (Supplementary Fig. S16).

**T Cell Functional Experiments:** Proliferation and cytotoxicity were evaluated as previously described^1,2^. Briefly, to determine proliferative ability, CART cells were plated at a 1:1 effector-to-target cell ratio on Day 0 of the proliferation assay. On Day 3, 100μL of media was removed from each well and cryopreserved for cytokine release experiments as described in a subsequent section. Cells were then fed with 100μL of T cell media and incubated until Day 5. On Day 5, the absolute CD3^+^ cell count was determined via flow cytometry with CD3 (clone SK7) APC-Cy7 (cat #344818, BioLegend, San Diego, CA, USA). For proliferation assays that tested the effects of hrIL-4 or IL-4 mAb, the 100μL of media added on Day 3 contained 20ng/mL hrIL-4 (cat #78045, STEMCELL Technologies, Vancouver, Canada), 10μg/mL *InVivo*MAb anti-human *IL*-4 (cat #BE0240, BioXCell, Lebanon, NH, USA), or 10μg/mL *InVivo*MAb rat IgG1 isotype control, anti-horseradish peroxidase (cat #BE0088, BioXCell, Lebanon, NH, USA). Briefly for cytotoxicity assays, luciferase^+^ target cells (JeKo-1) were incubated at the indicated effector-to-target cell ratios for 48 hours as listed in the specific experiment. Killing was calculated by bioluminescence imaging on a GloMax Explorer (Promega, Madison, WI, USA) after treating samples with 1μL D-luciferin (30μg/mL) per 100μL sample volume (Gold Biotechnology, St. Louis, MO, USA), for 5 minutes prior to imaging.

**RNA Isolation and Reverse Transcription-Quantitative Polymerase Chain Reaction (RT-qPCR) for EOMES:** Total RNA was extracted with QIAzol lysis reagent (Qiagen, Gaithersburg, MD, USA), RNeasy Plus Mini Kit (Qiagen, Gaithersburg, MD, USA), and RNase-Free DNase Set (Qiagen, Gaithersburg, MD, USA) according to the manufacturer’s protocol. cDNA was generated using iScript Advanced cDNA Kit for RT-qPCR (Bio-Rad, Hercules, CA, USA) according to manufacturer’s instructions. RT-qPCR reaction was completed according to manufacturer’s instructions for RT-qPCR SsoAdvanced Universal SYBR Green Supermix (cat # S7563, Invitrogen, Carlsbad, CA, USA). The primer sequences used were as follows: forward primer EOMES (5’-GGCCTCTGTGGCTCAAATTC-3’), reverse primer EOMES (5’-GCAGTGGGATTGAGTCCGTT-3’), forward primer GAPDH (5’-GGAGCGAGATCCCTCCAAAAT-3’)^8^, reverse primer GAPDH (5’-GGCTGTTGTCATACTTCTCATGG-3’)^8^, forward primer TBP (5’-CTCACAGGTCAAAGGTTTAC-3’)^9^, reverse primer TBP (5’-GCTGAGGTTGCAGGAATTGA-3’)^9^.

**Mantle Cell Lymphoma Xenograft Mouse Model:** Male and female 6- to 8-week-old NOD-SCID-IL2rγ^-/-^ (NSG) mice were purchased from Jackson Laboratories and were cared for within the Department of Comparative Medicine at the Mayo Clinic under an approved Institutional Animal Care and Use Committee protocol (A00001767-16-R22). Mice were allowed to acclimate for two weeks before being included in experiments. In the mantle cell lymphoma xenograft mouse model which stress-tested CART19-28ζ cells, 1x10^6^ luciferase^+^ JeKo-1 cells were engrafted via tail vein injection. Tumor burden was monitored with bioluminescent imaging as previously described^1^. Once the tumor burden reached approximately 1x10^8^ photons/second, mice were weighed and randomized based on tumor burden into treatment groups. CART19-28ζ cells were administered via tail vein injection as indicated in each specific experiment. For the mice receiving a combination of CART19-28ζ cells and either an IL-4 mAb (MP4-25D2, cat #BE0240, BioXCell, Lebanon, NH, USA) or an IgG control antibody (cat #BE0088, BioXCell, Lebanon, NH, USA), mice received weekly intraperitoneal injections of 10mg/kg antibody.

Following the start of treatment, mice were followed for tumor burden as determined with bioluminescence imaging, CART cell expansion and cytokine profile of peripheral blood, and overall survival. Following peripheral blood sampling, 50μL of blood was lysed of red blood cells with FACS^TM^ Lysing Solution (cat #349202, BD Biosciences, San Diego, CA, USA) prior to antibody staining with anti-human CD45 (clone HI30) BV421 (cat #304032, BioLegend, San Diego, CA, USA), anti-mouse CD45 (clone 30-F11) APC-eFluor780 (cat #47-0451-82, eBioscience, San Diego, CA, USA), anti-human CD3 (clone SK7) BV605 (cat #344836, BioLegend, San Diego, CA, USA), anti-human CD20, anti-human CD8, anti-human PD-1, anti-human TIM-3, and anti-human CTLA-4. Flow cytometry was then performed to determine the absolute CD3^+^ T cell count and the expression of inhibitory receptors. Serum from peripheral blood was then used for multiplex analysis of cytokines as described below.

**Serum and Supernatant Analysis of Cytokine Concentration:** Cytokine concentration in the supernatant of *in vitro* assays and in the serum of mice was determined with MILLIPLEX MAP Human High Sensitivity T Cell Panel Premixed 13-plex (cat #HSTCMAG28PMX13BK, Millipore Sigma, Ontario, Canada) according to the kit’s instructions with 25μL of either serum or supernatant per sample. Analysis was completed with Belysa software after running the samples on a Luminex 200 (Millipore Sigma, Ontario, Canada).

**RNA Sequencing:** For RNA sequencing of healthy donor CART cells, CART cells were first isolated from co-culture using combined CD4 and CD8 microbeads (cat #130-045-101 and cat #130-045-201, Miltenyi Biotec, Auburn, CA, USA) as described above. Then, RNA was isolated from 1x10^6^ cells by using the miRNeasy Micro kit (Qiagen, Gaithersburg, MD, USA) and treated with RNase-Free DNase Set (Qiagen, Gaithersburg, MD, USA). RNA quality was assessed by High Sensitivity RNA Tapestation (Agilent Technologies Inc., California, USA) before library construction with the SMARTer Stranded Total RNA-Seq Kit v2 – Pico Input Mammalian (Takara Bio USA INC., California, USA). Next, RNA sequencing was performed on an Illumina NovaSeq S4 (Illumina, California, USA). CD Genomics provided the output as fastq files through a file transfer protocol (ftp). Fastq files were first evaluated for quality using FASTQC. Then, Cutadapt was used to remove Illumina adapters before FASTQC was run again to evaluate quality after adapter removal. Next, the paired end reads for each condition were aligned with STAR using the genome reference consortium human build 38 patch release 13 (GRCh38.p13) downloaded from NCBI^10^. HTSeq was used to generate expression counts for each gene, and DESeq2 was used to normalize the data and calculate differential expression^11,12^. Differentially expressed genes were determined based on a false discovery rate (FDR) less than 0.05. Volcano plots were generated with the package EnhancedVolcano^13^. Heatmaps were generated with the R package pheatmap. Pathway analysis was conducted through the use of Qiagen IPA (Qiagen Inc., <https://digitalinsights.qiagen.com/IPA>)^14^.

RNA sequencing of pre-infusion axi-cel samples was completed by Kite Pharma and the sequencing files were shared with the Kenderian Laboratory. Briefly, frozen cell pellets were used for RNA isolation and library preparation. Then, strand-specific RNA sequencing (2x150 bp) with Poly-A selected RNA was performed. Following sequencing, the latest human genome (GRCh38.84) was downloaded from the Ensembl database and STAR was used to align the paired end reads for each sample to the genome. HTSeq was used to generate expression counts for each gene, and DESeq2 was used to normalize the data and calculate differential expression^11,12^. Differential expression was determined based on a p-value less than 0.05. Volcano plots were generated with the package EnhancedVolcano^13^. Heatmaps were generated with the R package pheatmap. Pathway analysis was conducted through the use of Qiagen IPA (Qiagen Inc., <https://digitalinsights.qiagen.com/IPA>)^14-16^.

**ATAC Sequencing:** For all samples, 1x10^5^ cells were washed in PBS and pelleted. The cell pellet was resuspended in 100μL freezing medium, composed of 10% dimethyl sulfoxide (Sigma, St. Louis, MO, USA) and 90% fetal bovine serum (FBS, Sigma, St. Louis, MO, USA). Then, it was stored at -80°C prior to shipment to CD Genomics for library preparation with the Nextera kit and sequencing services with the HISEQ 4000. CD Genomics provided the output as fastq files through ftp. Quality check was performed first with FASTQC. Then, the Nextera adapter sequences were trimmed and a minimum length of 45 base-pairs was employed per ENCODE recommendations using Cutadapt. FASTQC was then re-run to ensure quality. Next, paired-end reads for each sample were aligned to the UCSC reference genome for human genome 38 (hg38), patch release 13 with Bowtie2. The resulting bam file was sorted with samtools and mitochondrial reads were removed with the BAMQC program. Peak calling was performed on the sorted and cleaned bam file using the MACS2 package with broad peak calling. Differential peak accessibility analysis was conducted with DESeq2 in the DiffBind package after removing the blacklisted regions associated with hg38 and normalizing the data based on library size. Next, the differentially accessible peaks were annotated using the R package “ChIPSeeker”^17^. Differentially accessible genes were determined with using FDR less than 0.05 for healthy donor CART samples and with p-value less than 0.05 for patient CART samples. Motif analysis was conducted using MEME suites. Chromatin accessibility profiles were created with the UCSC genome browser with averaged BigWig files for each condition. Briefly, Bigwig files were created from sorted BAM files using the bamCoverage function. Then, averaged BigWig files for each experimental condition were generated with the mean function of Wiggletools.

**CRISPR Screen Considerations:** The protocol used for this CRISPR screen was adapted from a previously published article^18^. Briefly, the library A of the Human CRISPR Knockout Pooled Library (GeCKO v2) (cat #1000000048, AddGene, Cambridge, MA, USA) was amplified through the use of Endura Electrocompetent Cells (cat # 60242-1, Biosearch Technologies, Middlesex, UK) according to the previously described protocol^18^. An even sgRNA distribution of the amplified library was verified through next-generation sequencing and the use of MAGeCK analysis based on the calculated Gini index and the number of gRNAs in the library with zero counts^19^.

After verifying a low Gini index and number of unrepresented gRNAs in the amplified library, lentivirus was generated as previously described for CAR lentivirus^1^. Lentivirus for the library was titered using a puromycin selection method. On Day 2 of the CRISPR screen, CART cells were transduced with library lentivirus at an MOI of 0.3. From Days 3-8 of the screen, puromycin selection was conducted to purify the population of CART cells successfully transduced with the GeCKO library A lentivirus. Then, on Days 8 and 22 of the screen, CART cells were isolated from co-culture using combined CD4 and CD8 microbeads (cat #130-045-101 and cat #130-045-201, Miltenyi Biotec, Auburn, CA, USA) as described above and a pellet of 3.5x10^7^ cells CART cells were cryopreserved.

**CRISPR Screen (NGS Data Processing):** To prepare for sequencing, 3.5x10^7^ cells from baseline (Day 8) and chronically stimulated (Day 22) timepoints were washed and pelleted before storage at -20°C. Next, genomic deoxyribonucleic acid (gDNA) was isolated from each cell pellet with the Quick-DNA Midiprep Plus Kit (cat #D4075, Zymo Research, Irvine, CA, USA) according to the manufacturer’s protocol. An ethanol precipitation protocol was performed to enhance the purity of the gDNA following isolation. Next, the gDNA was prepared for sequencing through library PCR. In summary, the NEBNext® High-Fidelity 2X PCR Master Mix (cat #M0541S, New England BioLabs, Ipswich, MA, USA) was used to amplify the gRNA in the samples with the forward and reverse primers and the cycling conditions that were previously described in the referenced protocol paper^18^. Finally, the PCR reactions for each sample were pooled and the product was purified by running it on a 2% (wt/vol) agarose gel before extracting it with the QIAquick Gel Extraction Kit (cat #28704, Qiagen, Gaithersburg, MD, US).

Next, amplicon deep sequencing was performed by CD Genomics using the PE150 (Illumina, San Diego, CA, USA). CD Genomics provided the output as paired end fastq files through ftp. Paired-end read files for each sample were merged with bbmerge before using MAGeCK-VISPR to perform quality and differential expression analysis^19^. A volcano plot depicting the positively and negatively selected gRNAs was generated with ggplot and gene set enrichment analysis was performed on both the top negatively and the top positively-selected genes (FDR< 0.25) to identify the top affected pathways using Enrichr and QIAGEN IPA.

**Statistics:** Prism Graph Pad (LA Jolla, CA, USA) and Microsoft Excel (Redmond, WA, USA) were used to analyze the experimental data. RT-qPCR results were analyzed using REST-MCS© version 2^20^.

**Tables:**

Table S1: Antibodies used for flow cytometry experiments in this study.
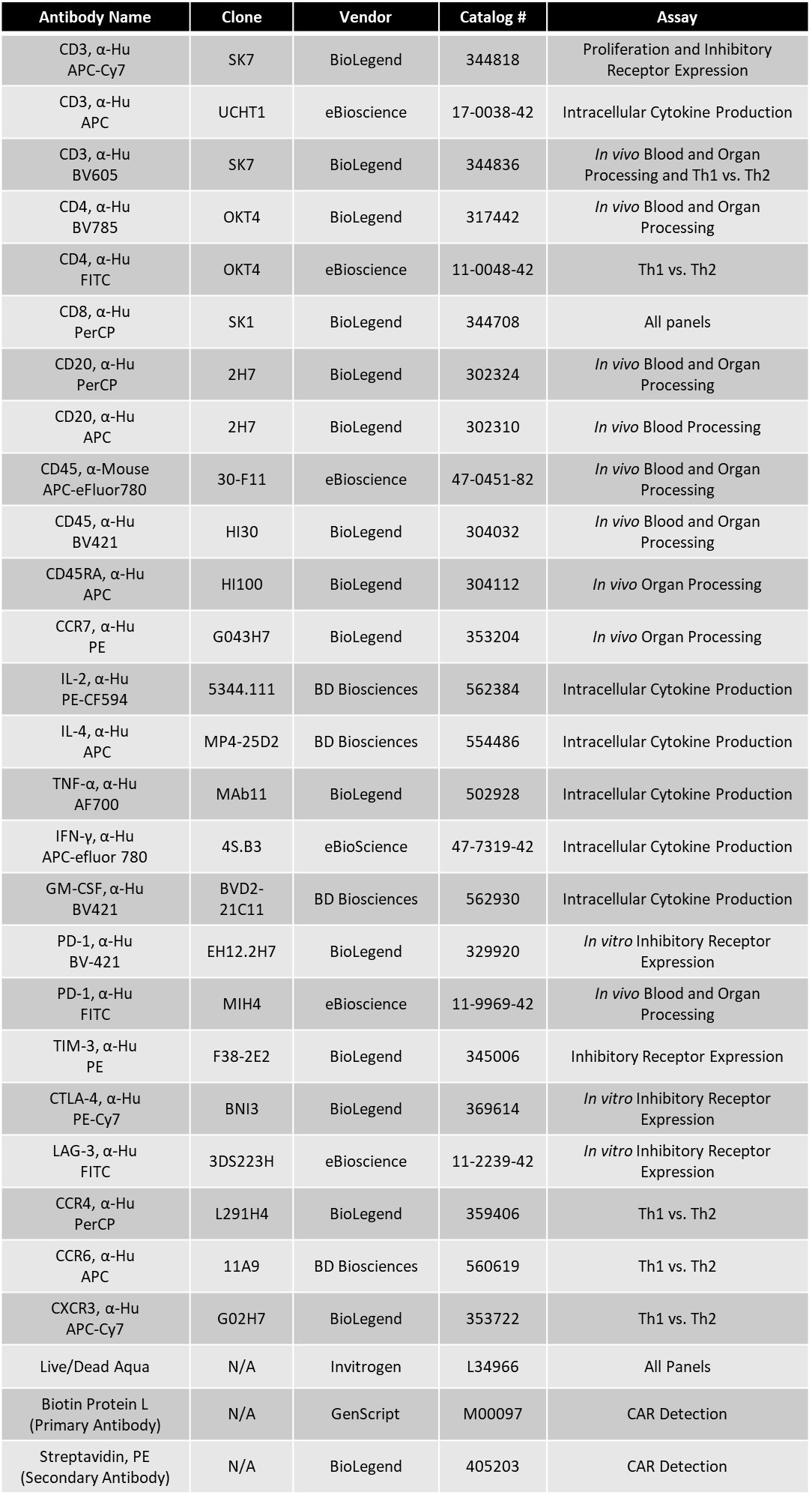

**Supplementary Figures:**

**supplementary Figure S1.**

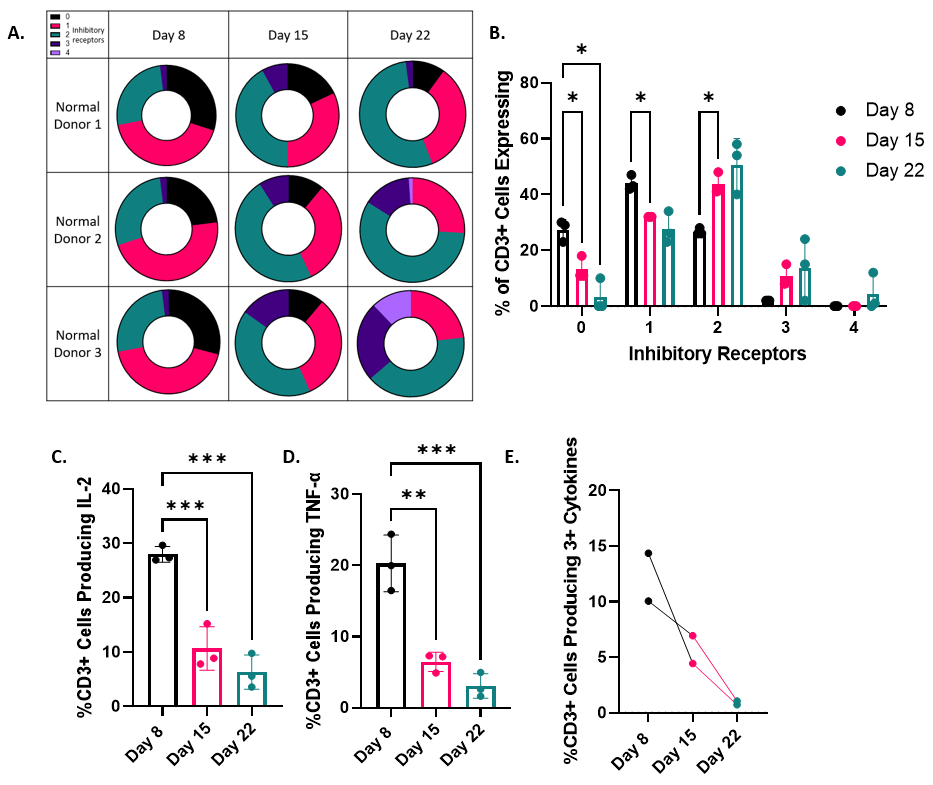

**Supplementary Figure S1.** CART19-28ζ cells chronically stimulated by JeKo-1 target cells using the *in vitro* model for exhaustion show phenotypical and functional signs of exhaustion. **A.** Circle plot showing the percent of CART cells expressing multiple inhibitory receptors (0 – black, 1 – pink, 2- green, 3 – dark purple, 4 – light purple) over time as determined with flow cytometric detection of CD3^+^ cells positive for PD-1, TIM-3, CTLA-4, and/or LAG-3 on Days 8, 15, and 22 of the *in vitro* model for exhaustion where CART19-28ζ cells were chronically stimulated with JeKo-1 target cells. **B.** Bar graph quantifying the circle plots in A. (Two-way ANOVA with three biological replicates, two technical replicates per biologic replicate). **C-D.** The percent of CART19-28ζ cells producing the effector cytokines IL-2 and TNF-α, respectively, on Day 8, 15, and 22. This was determined by strongly stimulating CART cells at a 1:5 E:T cell ratio for four hours before performing intracellular staining for the cytokines (One-way ANOVA with three biological replicates, two technical replicates per biologic replicate). **E.** The percent of CART19-28ζ cells producing three or more cytokines after strongly stimulating them at a 1:5 E:T cell ratio on Days 8, 15, and 22 of the *in vitro* model for exhaustion before performing intracellular staining for the following cytokines: IL-2, TNF-α, IFN-γ, and GM-CSF (Data from two biological replicates, two technical replicates per biological replicate). (*p<0.05, **p<0.01, ***p<0.001)

**supplementary Figure S2.**

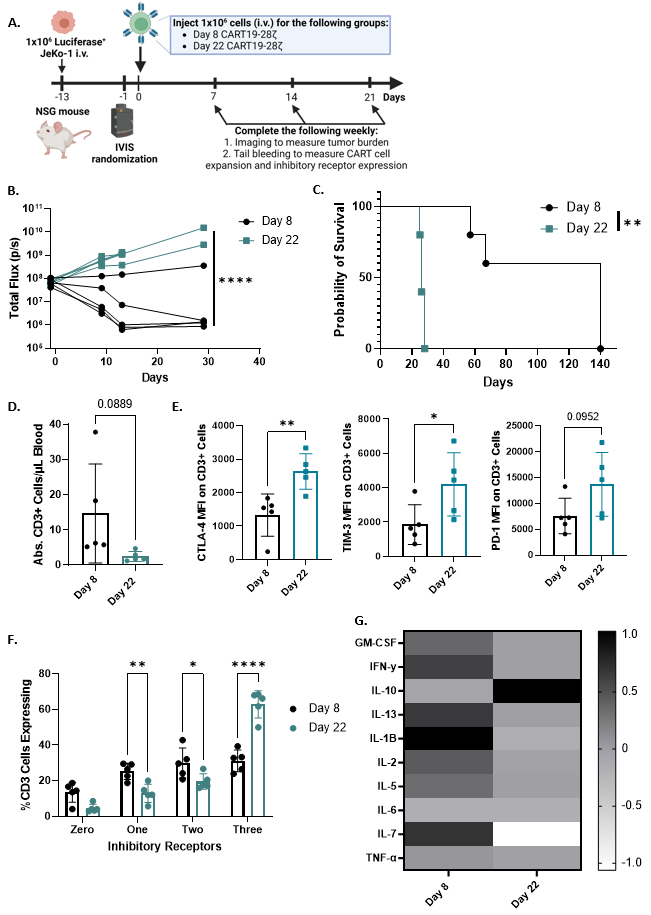

**Supplementary Figure S2.** Day 22 CART19-28ζ cells from the *in vitro* model for exhaustion showed reduced overall efficacy in two separate mantle cell lymphoma xenograft mouse models with different T cell donors. **A.** Schematic depicting an *in vivo* mantle cell lymphoma xenograft mouse model to induce stress in CART19 cells. **B.** Tumor burden overtime in a mantle cell lymphoma xenograft mouse model as determined by bioluminescence imaging of the luciferase^+^ tumor (Two-way ANOVA with n=5 mice per group, results from replicate experiment). **C.** Overall survival curve of mice in a mantle cell lymphoma xenograft mouse model (Log-rank (Mantle-Cox) test with n=5 mice per group, results from replicate experiment). **D.** Absolute human CD3^+^ cells per µL of peripheral blood as determined by flow cytometry on Day 14 of a mantle cell lymphoma xenograft mouse model (T-test with n=5 mice per group, results from replicate experiment). **E.** Mean fluorescence intensity of the inhibitory receptors CTLA-4, TIM-3, and PD-1 on human CD3^+^ cells in the peripheral blood of mice treated with either Day 8 or Day 22 CART19-28ζ cells on Day 14 of a mantle cell lymphoma xenograft mouse model (Two-way ANOVA with n=5 mice per group, results from replicate experiment). **F.** The percent of CART19-28ζ cells expressing multiple inhibitory receptors on Day 15 of a mantle cell lymphoma xenograft mouse model. This was determined through tail vein bleeding and flow cytometric detection of human CD3^+^ cells that are positive for PD-1, TIM-3, and/or CTLA-4 (Two-way ANOVA with n=5 mice per group). **G.** Heat map showing median of normalized values for cytokine concentration from each group of mice in the mantle cell lymphoma xenograft mouse model treated with either Day 8 or Day 22 CART19-28ζ cells (n=5 mice per group). (*p<0.05, **p<0.01, ****p<0.0001)

**Supplementary Figure S3.**

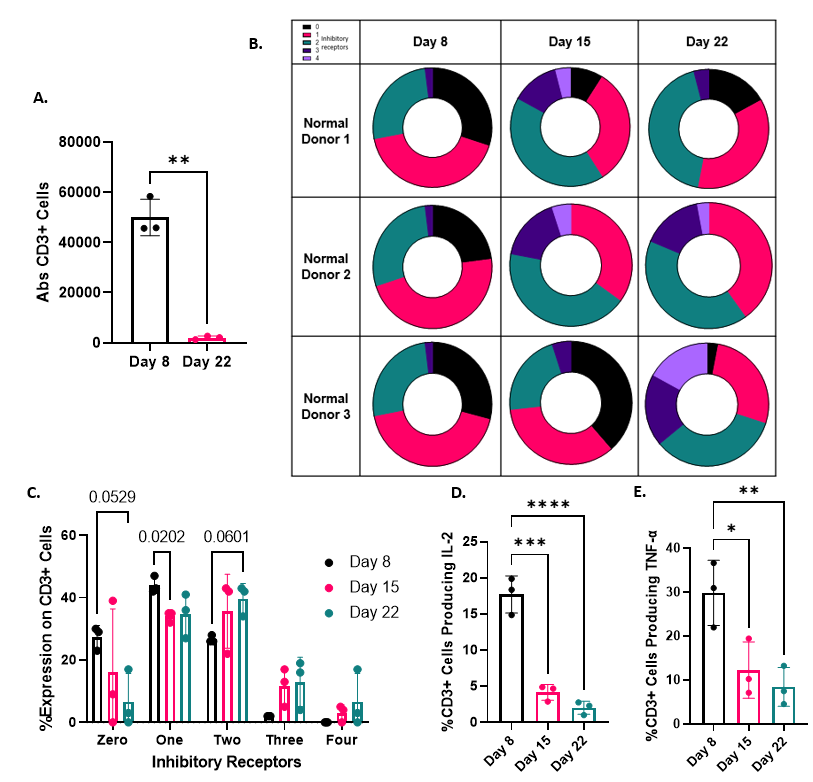

**Supplementary Figure S3.** Chronic stimulation of CART19-28ζ cells with the CD19^+^ acute lymphoblastic leukemia (ALL) cell line, NALM6, using the *in vitro* model for exhaustion results in phenotypical and functional signs of exhaustion. **A.** The absolute count of CD3^+^ T cells as determined with flow cytometry following co-culture of either Day 8 or Day 22 CART19-28ζ cells with NALM6 target cells at a 1:1 E:T cell ratio for 5-days (t-test with three biological replicates, two technical replicates per biologic replicate). **B.** Circle plot showing the percent of CART cells expressing multiple inhibitory receptors (0 – black, 1 – pink, 2- green, 3 – dark purple, 4 – light purple) over time as determined with flow cytometric detection of CD3^+^ cells positive for PD-1, TIM-3, CTLA-4, and/or LAG-3 on Days 8, 15, and 22 of the *in vitro* model for exhaustion where CART19-28ζ cells were chronically stimulated with NALM6 target cells. **C.** Bar graph quantifying the circle plots in **B.** (Two-way ANOVA with three biological replicates, two technical replicates per biological replicate). **D-E.** The percent of CD3^+^ cells producing IL-2 and TNF-α as determined with intracellular staining and flow cytometry after either Day 8, Day 15, or Day 22 CART19-28ζ cells were co-cultured with NALM-6 cells at a 1:5 E:T cell ratio for four hours (One-way ANOVA with three biological replicates, two technical replicates per biological replicate). (*p<0.05, **p<0.01, ***p<0.001, ****p<0.0001)

**Supplementary Figure S4.**

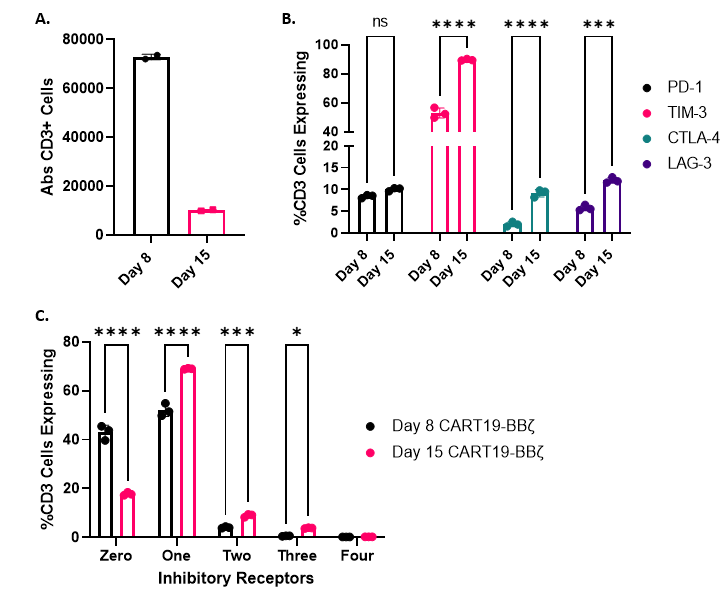

**Supplementary Figure S4.** Chronic stimulation of CART19-BBζ cells with CD19^+^ JeKo-1 cells using the *in vitro* model for exhaustion results in phenotypic and functional signs of exhaustion. **A.** Absolute CD3^+^ count as determined with flow cytometry after culturing either Day 8 or Day 15 CART19-BBζ cells with JeKo-1 cells for 5-days at a 1:1 E:T cell ratio (t-test with one biological replicate and two technical replicates). **B.** The percent of CART19-BBζ cells positive for either PD-1, TIM-3, CTLA-4, or LAG-3 as determined by flow cytometry on Day 8 and Day 15 of the *in vitro* model for exhaustion (Two-way ANOVA with one biological replicate and three technical replicates). **C.** The percent of CART19-BBζ cells positive for either 0, 1, 2, 3, or 4 inhibitory receptors as determined with flow cytometry on Day 8 and Day 15 of the *in vitro* model for exhaustion after staining for PD-1, TIM-3, CTLA-4, and LAG-3 (Two-way ANOVA with one biological replicate and three technical replicates). (*p<0.05, ***p<0.001, ****p<0.0001, ns: p>0.05)

**Supplementary Figure S5.**

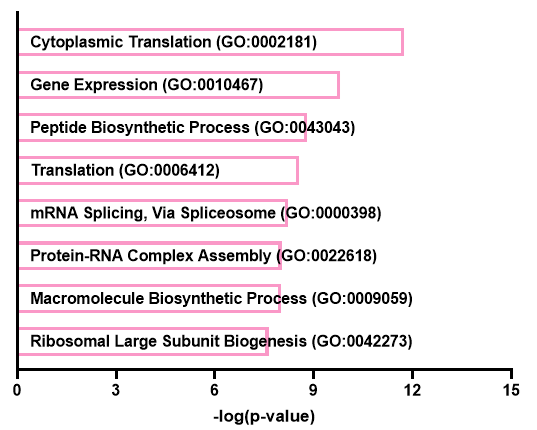

**Supplementary Figure S5.** Gene set enrichment analysis of negatively selected genes (FDR<0.25) in the genome-wide CRISPR knockout screen.

**Supplementary Figure S6.**

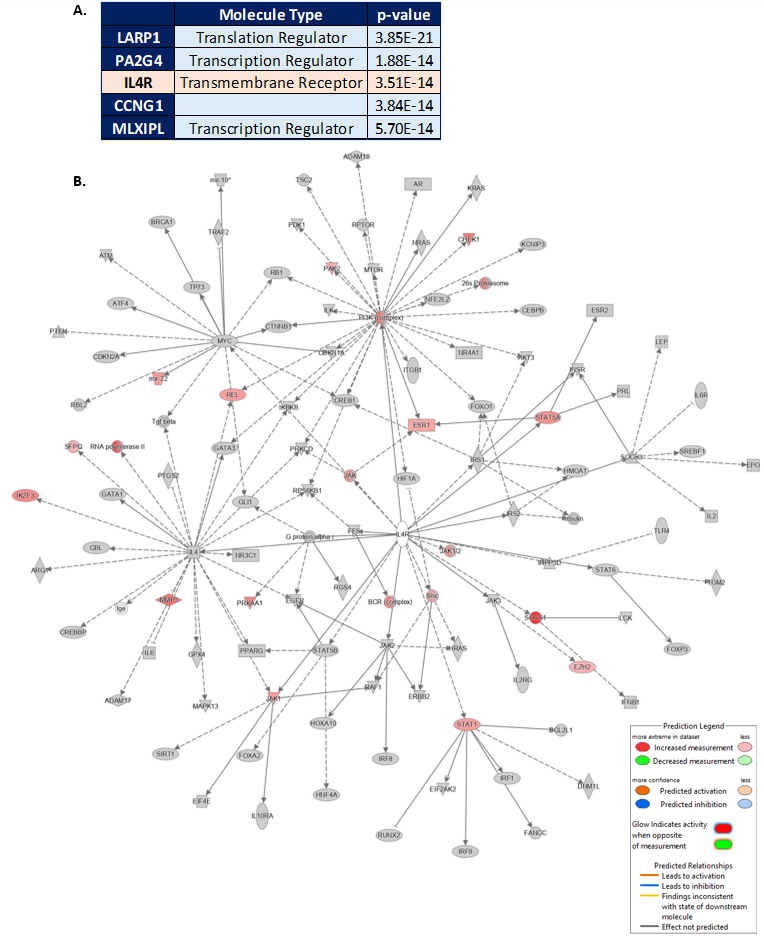

**Supplementary Figure S6.** Ingenuity pathway analysis of positively selected genes in the genome-wide CRISPR knockout screen. **A.** Top five casual networks identified through analysis of genes that were positively selected for by Day 22 of the genome-wide CRISPR knockout screen. **B.** QIAGEN IPA network for IL4R and affected genes as identified in the list of positively selected genes. (Positive selection in the CRISPR screen was defined as FDR<0.25)

**Supplementary Figure S7.**

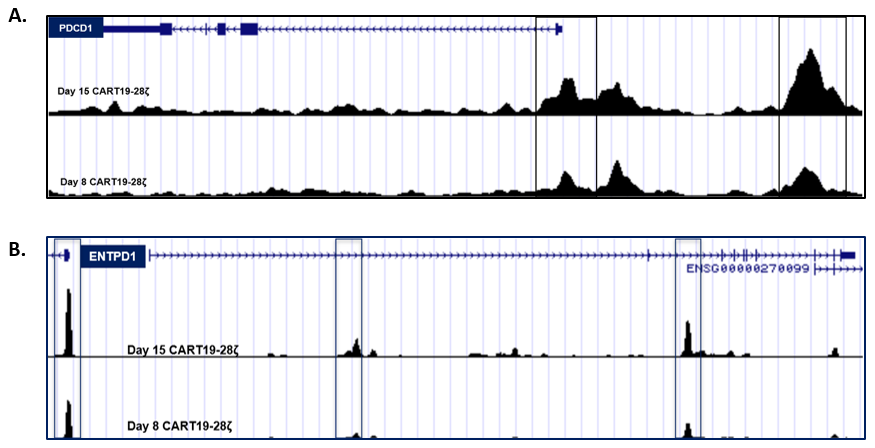

**Supplementary Figure S7.** Transcriptomic and chromatin accessibility interrogation of baseline and chronically stimulated CART19-28ζ cells from the *in vitro* model for exhaustion reveal key epigenetic signatures of exhaustion. **A-B.** ATAC signal track of selected gene loci (*PDCD1* and *ENTPD1*) showing averaged signal for the three biological replicates at each timepoint.

**Supplementary Figure S8.**

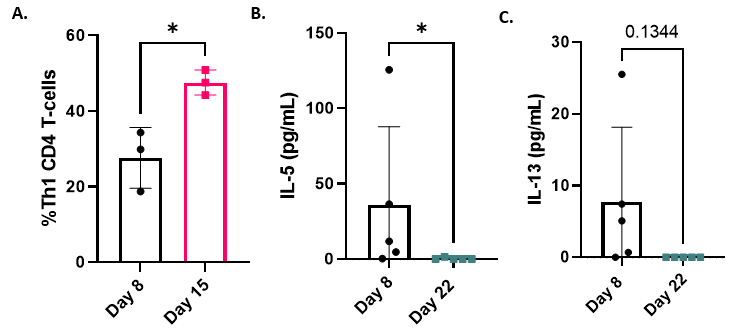

**Supplementary Figure S8.** Changes in the Th1/Th2 pathway during the *in vitro* model for exhaustion. **A.** The percent of Th1 CD4^+^ CART19-28ζ cells as defined by CCR6^-^CCR4^-^CXCR3^+^ cells by flow cytometry. **B-C.** The concentration of IL-5 and IL-13 in the serum by Multiplex assay of JeKo-1 xenograft mice two weeks after the injection of either Day 8 or Day 22 CART19-28ζ cells (n=5 mice per group, t-test). (*p<0.05)

**Supplementary Figure S9.**

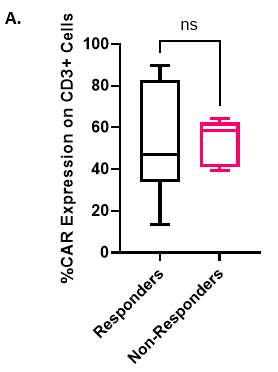

**Supplementary Figure S9.** No difference in CAR+ T cells between responders and non-responders to CART19-28ζ in the ZUMA-1 clinical trial **A.** The percent of CD3^+^ cells expressing CAR as determined by positive staining for an anti-Whitlow linker antibody using flow cytometry. (t-test with n=6 responders and n=6 non-responder samples, ns: p>0.05)

**Supplementary Figure S10.**

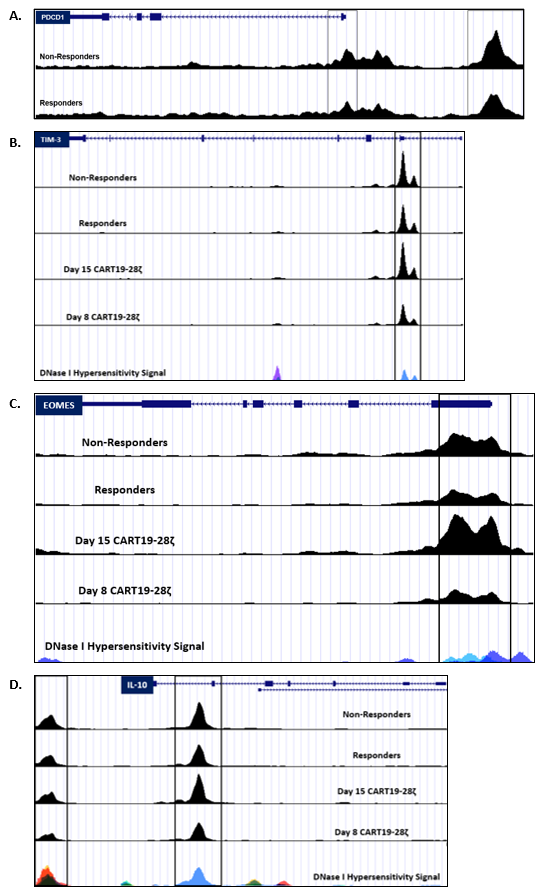

**Supplementary Figure S10.** Chromatin accessibility analysis of baseline axi-cel products from 6 responders and 6 non-responders in the ZUMA-1 clinical trial **A-D.** ATAC signal track of selected, exhaustion-related gene loci (*PDCD1, HAVCR2* (TIM-3), *EOMES, IL-10*) based on averaged signal for the biological replicates for each condition (n=6 for responders and n=6 for non-responders; n=3 for Day 15 and n= 3 Day 8 CART29-28ζ).

**Supplementary Figure S11.**

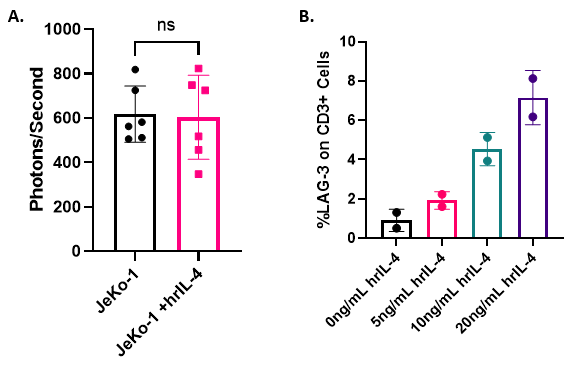

**Supplementary Figure S11.** The effect of treatment with human recombinant IL-4 (hrIL-4) *in vitro.* **A.** The luminescence (photons/second) of luciferase^+^ JeKo-1 cells after 48 hours of treatment with wither diluent or 20ng/mL hrIL-4 (t-test, ns: p>0.05, six technical replicates). **B.** The percent of CART19-28ζ cells expressing the inhibitory receptor LAG-3 after one week of chronic stimulation with JeKo-1 cells in the presence of either 0, 5, 10, or 20ng/mL hrIL-4 (two technical replicates).

**Supplementary Figure S12.**

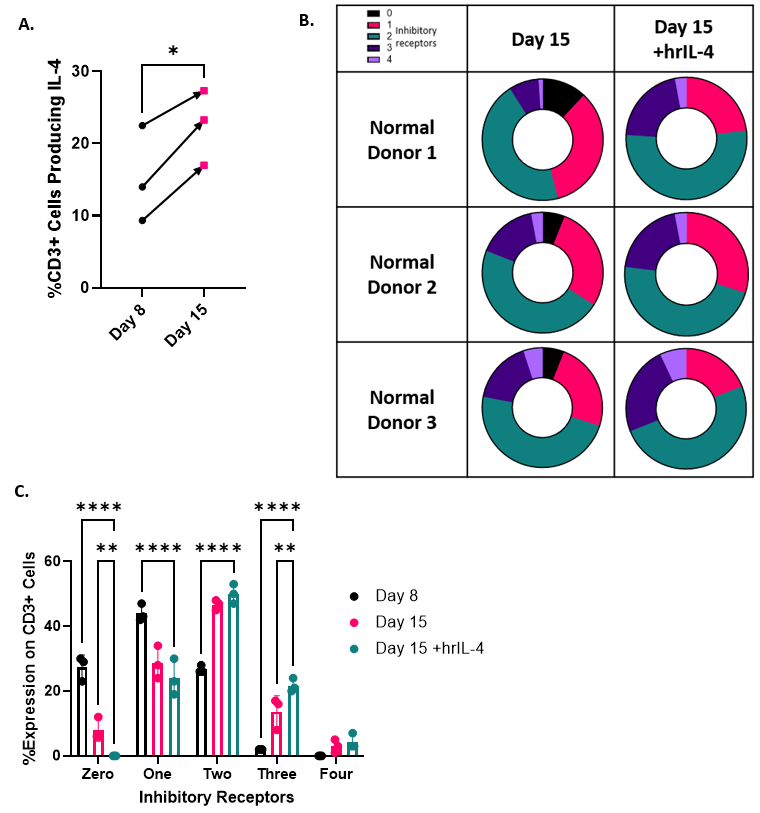

**Supplementary Figure S12.** Changes related to IL-4 are independent of stimulation condition of the CART cells. **A.** Change in the percent of CART19-28ζ cells producing IL-4 following chronic stimulation with irradiated JeKo-1 from Day 8 to Day 15. Production of IL-4 was determined with intracellular staining for flow cytometry following 4 hours of co-culture with either Day 8 or Day 15 CART19-28ζ cells and JeKo-1 cells at a 1:5 E:T cell ratio (t-test with three biological replicates, two technical replicates per biological replicate). **B-C.** Circle and bar graphs showing the change in the percent of CART19-28ζ cells expressing 0, 1, 2, 3, or 4 inhibitory receptors by flow cytometry from Day 8 to Day 15 CART19-28ζ cells that were chronically stimulated by irradiated JeKo-1 cells either in the presence of diluent or 20ng/mL hrIL-4 (Two-way ANOVA with three biological replicates, two technical replicates per biological replicate). (*p<0.05, **p<0.01, ****p<0.0001)

**Supplementary Figure S13.**

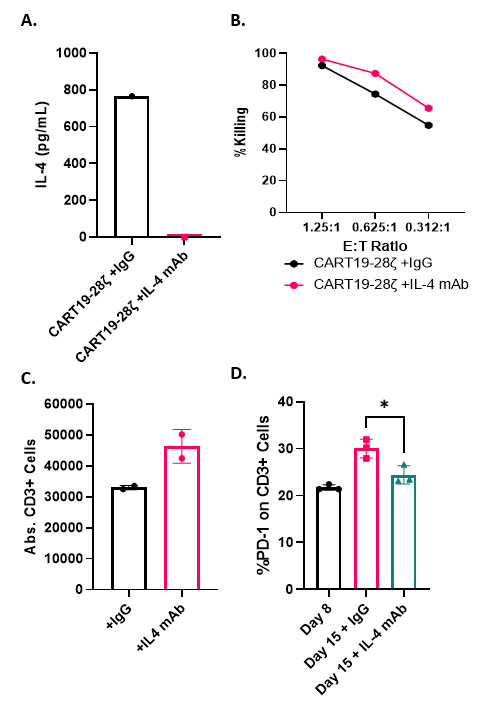

**Supplementary Figure S13.** The effect of IL-4 neutralization with a monoclonal antibody on CART19-28ζ function *in vitro.* **A.** The concentration of IL-4 in the supernatant after CART19-28ζ cells were co-cultured with JeKo-1 target cells at a 1:1 E:T ratio for three days in the presence of either 10μg/mL IL-4 mAb or IgG control antibody (one biological replicate). **B.** Cytotoxicity of CART19-28ζ cells that were co-cultured with luciferase^+^ JeKo-1 targets cells at various E:T ratios in the presence of either 10μg/mL IL-4 mAb or IgG control antibody for 48 hours (one biological replicate and two technical replicates). **C.** Absolute CD3^+^ cell count as determined by flow cytometry after culturing CART19-28ζ cells with JeKo-1 target cells at a 1:1 E:T cell ratio for 5-days in the presence of either 10μg/mL IL-4 mAb or IgG control antibody (one biological replicate and two technical replicates). **D.** The percent of CD3^+^ cells that are positive for PD-1 at Day 8 and Day 15 after being chronically stimulated with JeKo-1 cells in the presence of either 10μg/mL IL-4 mAb or IgG control antibody from Day 8 to Day 15 (One-way ANOVA with three biological replicates, two technical replicates per biological replicate, *p<0.05)

**Supplementary Figure S14.**

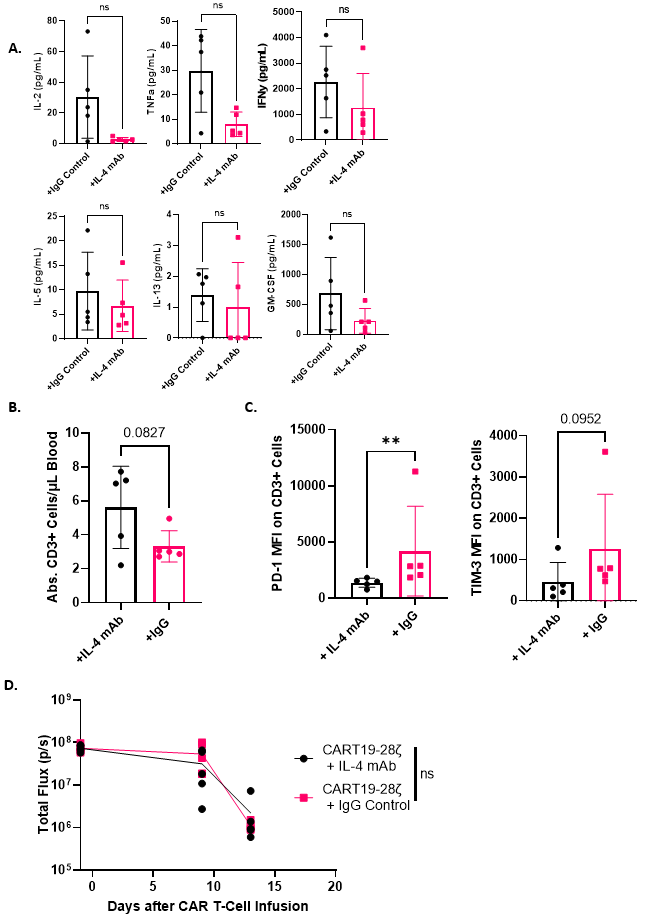

**Supplementary Figure S14.** The effect of IL-4 neutralization on CART19-28ζ efficacy in a stress and low tumor burden mantle cell lymphoma xenograft mouse model. **A.** Individual plots showing the concentration of detected cytokines on week two of the mantle cell lymphoma xenograft stress mouse model treated with Day 8 CART19-28ζ cells in combination with either 10mg/kg IL-4 mAb or IgG control antibody. (t-test, n=5 mice per group). **B.** CART cell expansion *in vivo* as determined by absolute CD3^+^ count by flow cytometry per μL of blood two weeks following CART cell infusion in a low tumor burden mantle cell lymphoma xenograft mouse model (t-test, n=5 mice per group). **C.** The mean fluorescence intensity of the inhibitory receptors PD-1 and TIM-3 by flow cytometry on CART cells in mice treated with either CART19-28ζ cells and 10mg/kg IL-4 mAb or CART19-28ζ cells and 10mg/kg IgG control antibody on week two of a low tumor burden mantle cell lymphoma xenograft mouse model (Two-way ANOVA with n=5 mice per group). **D.** Tumor progression as measured with bioluminescence imaging of luciferase^+^ JeKo-1 cells in a low tumor burden mantle cell lymphoma xenograft mouse model where the mice were treated with CART19-28ζ cells in combination with either 10mg/kg IL-4 mAb or IgG control antibody (Two-way ANOVA with n=5 mice per group). (ns: p>0.05, ***p<0.001)

**Supplementary Figure S15.**

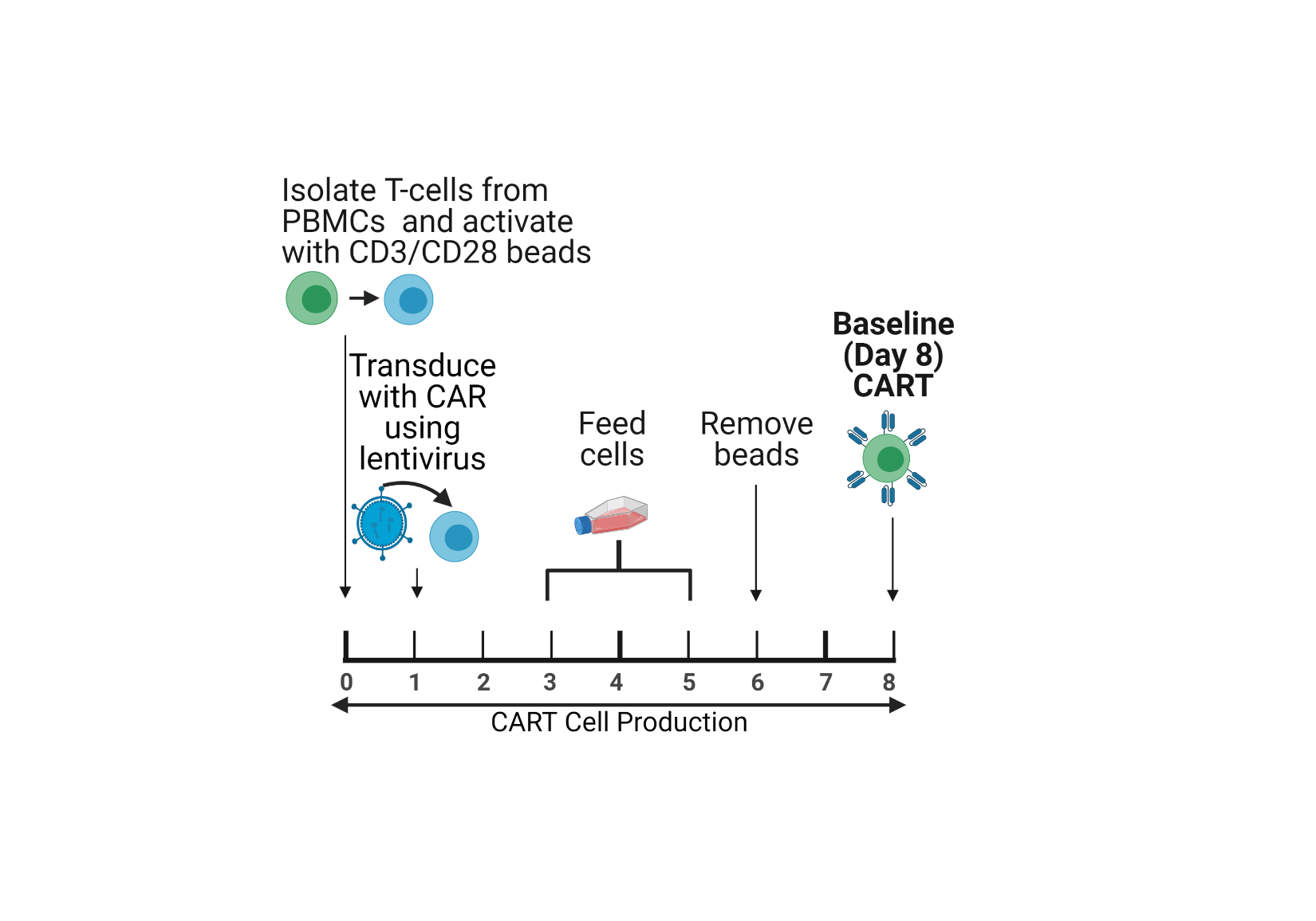

**Supplementary Figure S15.** Healthy donor CART cell production for *in vitro* and *in vivo* studies.

**Supplementary Figure S16.**

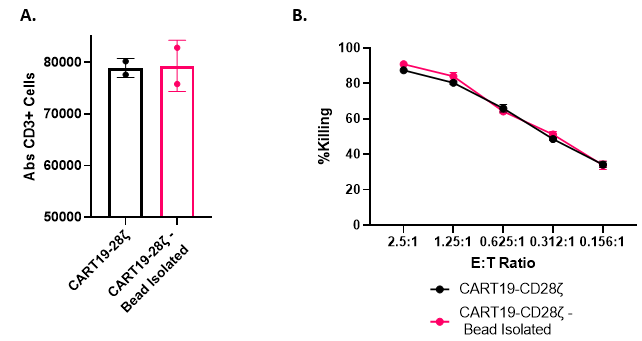

**Supplementary Figure S16.** The use of CD4^+^ and CD8^+^ beads to separate CART cells from co-culture does not impact functionality. **A.** Absolute CD3^+^ cell count as determined by flow cytometry after culturing CART19-28ζ cells, that had either underwent bead separation or did not, with JeKo-1 target cells at a 1:1 E:T cell ratio for 5-days (One biological replicate and two technical replicates). **B.** Cytotoxicity of CART19-28ζ cells that either underwent bead separation or did not before culture with luciferase^+^ JeKo-1 target cells at various E:T ratios for 48 hours (One biological replicate and two technical replicates).
